# APP-CTFβ/C99 oligomers drive synaptic vesicle tethering through C-terminal interactions

**DOI:** 10.64898/2026.06.25.734451

**Authors:** Akshay Kapadia, Elena Cjiffers, Stanley Van, Anne-Sophie Hafner

**Affiliations:** Donders Institute for Brain, Cognition and Behavior, Radboud University, Nijmegen, Netherlands; Interdisciplinary Institute for Neuroscience (IINS), Université de Bordeaux, UMR 5297 CNRS, 33076 Bordeaux, France

## Abstract

Proteolytic processing of the amyloid precursor protein (APP) generates a 99-amino acid precursor, β-carboxyl-terminal fragments (APP-CTFβ or C99). Upon γ-secretase inhibition, APP-CTFβ accumulates and induces synaptic defects, resulting in neuronal hyperactivity. However, mechanistic insights in the critical role of APP-CTFβ has not been completely elucidated. Here, we show that in primary neurons expressing human APP-CTFβ (C99) variants, acute γ-secretase inhibition selectively increases evoked synaptic vesicle release in cells, whereas deletion of the C-terminus abolishes this effect. Using single-molecule approaches and reconstituted membrane systems, we demonstrate that accumulation of APP-CTFβ promotes its oligomerisation. In particular, APP-CTFβ oligomers augment synaptic vesicle tethering via their C-terminal domain. This effect is driven by the interaction with synaptic vesicle proteins, independent of the YENPTY binding motif. Additionally, APP-CTFβ oligomers were associated with alterations in membrane lipid organization. Together, our findings identify APP-CTFβ oligomerization as a constitutional gain-of-function mechanism that enhances presynaptic vesicle tethering and release, providing mechanistic insight into how altered APP processing regulates synaptic activity.

**Short summary:** γ-Secretase-dependent accumulation of APP-CTFβ transforms a transient APP processing intermediate into a membrane-associated oligomeric scaffold that promotes synaptic vesicle tethering enhancing neurotransmitter release.

**Highlights:**

- APP-CTFβ self-assembles into higher-order oligomeric assemblies
- APP-CTFβ oligomers promote synaptic vesicle tethering via their C-terminal domain
- APP-CTFβ oligomerization drives a presynaptic gain-of-function linked to neuronal hyperactivity

## Introduction

The amyloid precursor protein (APP) is a type I transmembrane protein that undergoes sequential proteolytic processing by β- and γ-secretases, generating a range of fragments with distinct cellular functions (De Strooper & Annaert, 2000; Hoe *et al*, 2012). While extensive work has focused on the role of extracellular amyloid-β (Aβ) peptides in Alzheimer’s disease (AD) (Cai *et al*, 2023; Giorgio *et al*, 2024; Martinsson *et al*, 2022; Siddu *et al*, 2025; Takahashi *et al*, 2004; Wei *et al*, 2010; Wilson *et al*, 2025; Zott *et al*, 2019), increasing evidence suggests that intracellular APP-derived fragments also contribute to neuronal dysfunction (Chang & Suh, 2005; Daly *et al*, 1998; Kapadia *et al*, 2025; Kwart *et al*, 2019; Lauritzen *et al*, 2019b, 2012). In particular, β-secretase cleavage generates the membrane-bound β-carboxyl-terminal fragment APP-CTFβ (also termed C99), which is subsequently processed by γ-secretase. Under conditions where γ-secretase activity is impaired, APP-CTFβ accumulates within neurons and induces mitochondrial (Lee *et al*, 2022; Vaillant-Beuchot *et al*, 2021), endo-lysosomal (Lauritzen *et al*, 2019b; Bourgeois *et al*, 2018; Hung & Livesey, 2018; Kwart *et al*, 2019; Lauritzen *et al*, 2016, 2019a; Van Acker *et al*, 2019), proteasomal (Badot *et al*, 2026) and synaptic defects (Barthet *et al*, 2018; Burrinha *et al*, 2021; Daly *et al*, 1998; Essayan-Perez & Südhof, 2023; Ferrer-Raventós *et al*, 2023; Jordà-Siquier *et al*, 2022; Kapadia *et al*, 2025; Kwart *et al*, 2019; Luo *et al*, 2024; Martinsson *et al*, 2022; Schilling *et al*, 2023; Tambini *et al*, 2019; Yao *et al*, 2019; Zhou *et al*, 2022). Notably, oligomerization is a recurrent feature of APP-derived fragments, most prominently Aβ (Hayden & Teplow, 2013; Viola & Klein, 2015), yet whether analogous self-assembly of APP-CTFβ contributes to synaptic dysfunction remains largely unexplored. Several studies have further suggested that APP-CTFβ can self-associate (Pantelopulos *et al*, 2022, 2018; Badot *et al*, 2026; Perrin *et al*, 2020), although the functional consequences of these higher-order assemblies remain poorly understood. Whether such assemblies contribute directly to neuronal dysfunction or represent functionally active signalling states remains unknown.

Early stages of AD are characterized by increased neuronal excitability and network hyperactivity (Anastacio *et al*, 2022; Giorgio *et al*, 2024; Kazim *et al*, 2021; Ray *et al*, 2024; Samudra *et al*, 2023; Sciaccaluga *et al*, 2021; Tok *et al*, 2022). This phase precedes overt irreversible neurodegeneration and cognitive decline. Familial forms of early-onset AD, caused by mutations in *APP* and presenilin (*PSEN1* and *PSEN2*) genes, disrupt γ-secretase-mediated APP processing and alter the balance of APP-derived fragments (Annaert *et al*, 2000; Barthet *et al*, 2018; Chang & Suh, 2005; Kabir *et al*, 2020; Lauritzen *et al*, 2019b; Ray *et al*, 2024; Thordardottir *et al*, 2017). These changes are associated with synaptic dysfunction and elevated neuronal activity, suggesting that APP processing products, particularly APP-CTFβ, may directly influence presynaptic function (Barthet *et al*, 2018; Burrinha *et al*, 2021; Chang & Suh, 2005; Essayan-Perez & Südhof, 2023; Kapadia *et al*, 2025; Martinsson *et al*, 2022; Zhou *et al*, 2022; Luo *et al*, 2024). Consistent with this, the accumulation of APP-CTFβ is sufficient to enhance presynaptic glutamatergic activity, independent of extracellular Aβ (Das *et al*, 2021; Kapadia *et al*, 2025; Lauritzen *et al*, 2012; Tambini *et al*, 2019; Yao *et al*, 2019). Together, these observations suggest that APP-CTFβ is not merely a metabolic intermediate of APP processing, but an active regulator of presynaptic function capable of influencing neuronal excitability. Notably, APP-CTFβ is highly enriched at excitatory presynaptic terminals together with components of the amyloidogenic processing machinery (Kapadia *et al*, 2025; Das *et al*, 2021), placing it in a strategic position to directly influence synaptic activity.

At the presynaptic terminal, neurotransmitter release is governed by tightly coordinated processes. The molecular mechanisms controlling docking, priming and fusion have been extensively characterized (Dunant & Israël, 2000; Ermolyuk *et al*, 2012; Gormal & Meunier, 2022; Schmitt, 2024; Südhof, 2013, 2012). However, considerably less is known about how membrane-associated proteins, including APP and its proteolytic fragments, regulate earlier events such as vesicle tethering, vesicle availability, and local vesicle organization at presynaptic release sites. The intracellular C-terminal domain of APP has been implicated in endocytosis (Zadka *et al*, 2024) and intracellular signalling through interactions mediated by the canonical ‘YENPTY’ binding motif (Aow *et al*, 2023; Guénette *et al*, 2017; King & Scott Turner, 2004; Nhan *et al*, 2015; Ono *et al*, 1997; Zheng & Koo, 2006). Whether these interactions account for APP-CTFβ-dependent presynaptic effects remains unresolved.

Because APP-CTFβ remains membrane-anchored following β-secretase cleavage, its accumulation may influence both membrane organization and protein assemblies at presynaptic sites (Kapadia *et al*, 2025). Importantly, whether APP-CTFβ functions as a monomeric signalling intermediate or assembles into higher-order membrane-associated complexes that directly regulate presynaptic vesicle organization remains unknown.

To dissect the role of the APP-CTFβ and its intracellular domain, we compared wild-type APP and APP-CTFβ with variants that either disrupt the YENPTY interaction motif or remove the entire intracellular C-terminal region in primary neuronal and CHO cell-based model systems. Using neuronal imaging, single-molecule approaches, and reconstituted membrane systems, we demonstrate that APP-CTFβ oligomers act as membrane-associated assemblies that engage synaptic vesicle proteins and augment release probability. We show that γ-secretase-dependent accumulation of APP-CTFβ promotes its oligomerization, alters membrane lipid organization, and enhances synaptic vesicle tethering *via* its intracellular C-terminal domain. These findings identify a mechanism by which APP-CTFβ oligomers, through the intracellular C-terminal domain, directly modulate presynaptic vesicle tethering and local membrane organization. Taken together, these findings provide insights into how altered APP processing, particularly accumulation of APP-CTFβ, contributes to early synaptic dysfunction in AD.

## Results

### APP-CTFβ accumulates at excitatory presynaptic sites and enhances synaptic vesicle release following γ-secretase inhibition

To investigate how APP-CTFβ regulates synaptic function, we expressed human APP-FL and APP-CTFβ/C99 variants in rat primary neurons and examined their subcellular localization (Fig. 1A-G; S1) and functional impact on synaptic activity (Fig. 1H-M). We employed human APP-FL (695 wild-type isoform) which exhibits the presence of a large N-terminal extracellular ectodomain and is mainly processed *via* the α-secretase pathway, generating APP-CTFα (Parvathy *et al*, 1999; Thinakaran *et al*, 1996) alongside APP-CTFβ. Next, we compared the effects of APP-CTFβ (hereafter noted a C99^YENPTY^) with other C99 variants bearing targeted mutations at the ‘YENPTY’ binding domain (YENPTY→ANAETA), previously implicated in adaptor-mediated trafficking and intracellular signalling (C99^ANAETA^), as well as a C-terminal deletion mutant, completely lacking the intracellular C-terminal domain (C99^ΔCT^). Non-transfected cells (no DNA, only lipofectamine) are denoted as mock controls. Widely used Aβ17-24 sequence specific antibody clone: 4G8 was used in all experiments, as it uniformly recognises APP-FL and all C99 variants, regardless of the differences in their N- and C-terminal modifications. Immunofluorescence analyses revealed that APP-FL and all C99 variants were broadly distributed across neuronal compartments, including soma, dendrites, axons, and synapses, as defined by MAP2, neurofilament (NFL), and synapsin-1/2 markers (Fig. 1A-B), respectively. C99^YENPTY^ and C99^ANAETA^ displayed highly comparable subcellular distributions across neuronal compartments, indicating that mutation of the YENPTY motif does not substantially alter subcellular localization (Fig. 1A-B, S1E). The loss of the C-terminal domain in C99^ΔCT^ showed a relative enrichment within somato-dendritic compartments and reduced synaptic localization compared with the other C99 variants (Fig. S1E). Importantly, despite modest differences in subcellular localization, APP-FL and all C99 variants were readily detected in transfected neurons across all experimental conditions (Fig. 1A-B, S1C). This allowed us to use each construct as its own baseline and evaluate the effects of γ-secretase inhibition independently of expression differences between conditions.

**Figure 1.**
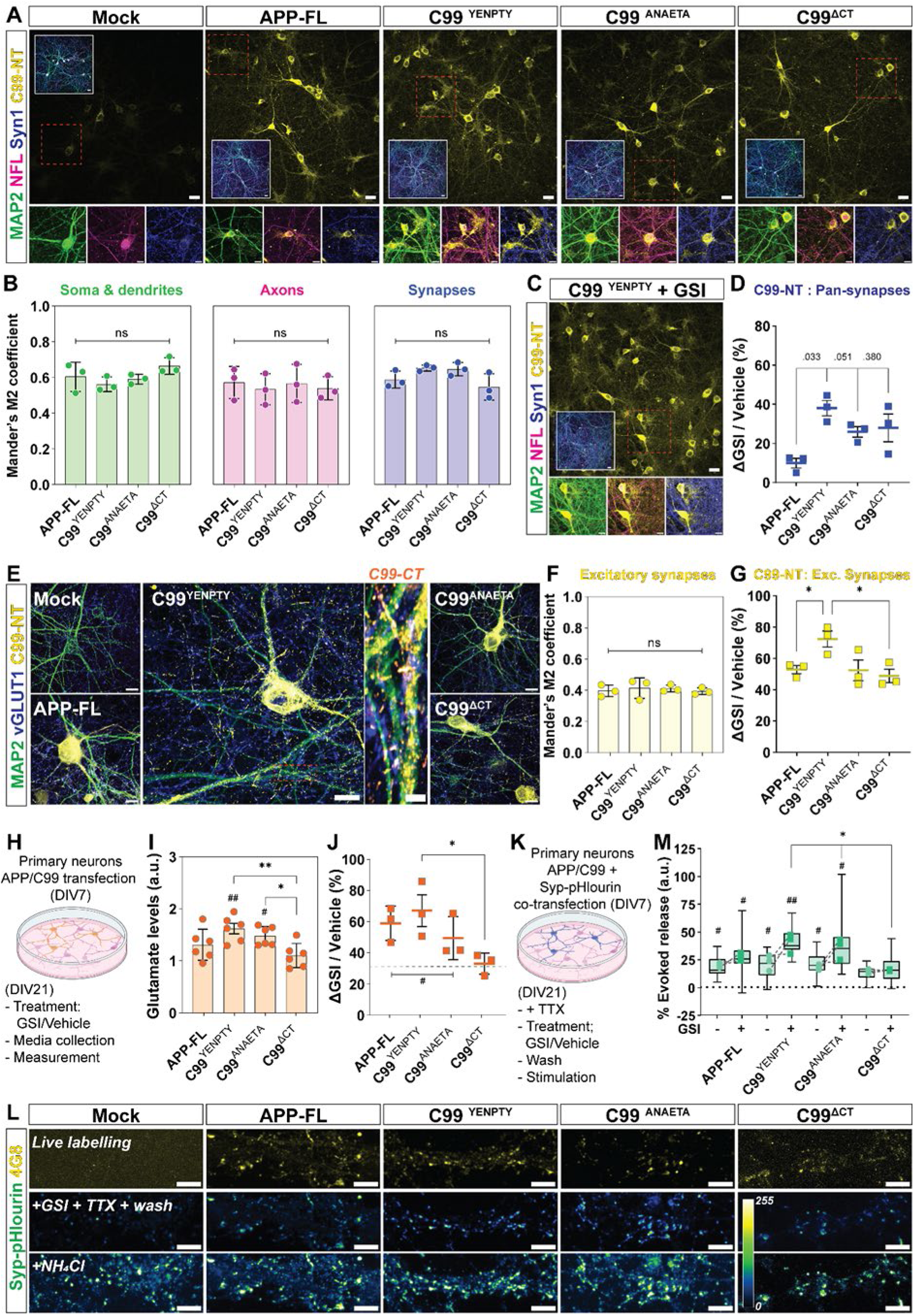
γ-Secretase inhibition alters subcellular localization of APP-CTFβ and enhances synaptic vesicle release. (A) Representative immunocytochemistry images of rat primary cortical neurons transfected with APP-FL or APP-CTFβ (C99) variants (C99^YENPTY^, C99^ANAETA^, C99^ΔCT^). APP and C99 species were detected using an Aβ mid-region-specific antibody (4G8, *yellow*) and co-stained for MAP2 (soma and dendrites, *green*), neurofilament light chain (NFL; axons, *magenta*), and synapsin-1/2 (Syn1/2; pan-synapses, *blue*). Non-transfected condition (lipofectamine without DNA) is indicated as mock. Lower panels show higher magnification views of neuronal processes and synaptic regions. Scale bars, 20 µm; zoom-in panels, 5 µm. **(B)** Bar plots depict Manders’ overlap coefficient (MOC-M2) of APP/C99 signals (4G8+) across somato-dendritic (MAP2+, *green*), axonal (NFL+, *magenta*), and synaptic (Syn1/2+, *blue*) compartments, under transient expression conditions. Individual points represent average from each independent experiment; bars, mean; error bars, S.D.; N = 3 independent transfected cultures. **(C)** Representative images depicting neurons expressing C99^YENPTY^ following γ-secretase inhibition (GSI; DAPT- 10 μM, 4 h) treatment. Scale bars, 20 µm; zoom-in panels, 5 µm. Additional images have been depicted in SI, Fig. S1D. **(D)** Scatter plot depicting percentage change (Δ) of APP/C99 overlap with Syn1/2 positive presynaptic boutons following GSI treatment relative to vehicle (untreated) controls. ΔGSI/Vehicle (%) = [(MOC_GSI_ - MOC_Vehicle_) / MOC_Vehicle_] × 100. Values are expressed as percentage change relative to the corresponding vehicle controls. Individual points represent average from each independent experiment; bold line, mean; error bars, S.D.; N = 3 independent transfected cultures. **(E)** Representative images showing co-localization of APP/C99 (4G8, *yellow*) with the excitatory synapse marker vesicular glutamatergic transporter 1 (vGLUT1, *blue*). Zoomed-in panels highlight co-distribution of C-terminal-specific (antibody clone: C1/6.1, C99-CT, *orange*) signals. Scale bars, 20 µm; zoom-in panels, 5 µm. **(F)** Bar plots depict MOC-M2 values of APP/C99 signals (4G8+) across vGLUT1+ excitatory presynaptic boutons, under transient expression conditions. Individual points represent average from each independent experiment; bars, mean; error bars, S.D.; N = 3 independent transfected cultures. **(G)** Scatter plot depicting percentage change (Δ) of APP/C99 overlap with vGLUT1 positive presynaptic boutons following GSI treatment relative to vehicle controls. Individual points represent average from each independent experiment; bold line, mean; error bars, S.D.; N = 3 independent transfected cultures. **(H)** Schematic of experimental design for glutamate release measurements performed using Promega Glutamate-Glo assay kit. **(I)** Bar plots showing extracellular glutamate levels measured from conditioned media of neurons expressing APP-FL or C99 variants under basal conditions and normalized to mock controls (dotted line). Individual points represent average from each independent experiment n = 6; bars, mean; error bars, S.D.; N = 3 independent transfected cultures. **(J)** Scatter plot depicting percentage change (Δ) of glutamate levels in extracellular glutamate levels following GSI treatment relative to vehicle controls, computed as ΔGSI/Vehicle (%) = [(GLUT_GSI_ - GLUT_Vehicle_) / GLUT_Vehicle_] × 100. Individual points represent average from each independent experiment; bold line, mean; error bars, S.D.; N = 3 independent transfected cultures. **(K)** Schematic of experimental workflow for synaptophysin-pHluorin (Syp-pHluorin) imaging in rat primary neurons. **(L)** Transfected neurons were treated with vehicle or GSI in the presence of TTX (1 μM, 4 h) and live-labelled with 4G8-biotin followed by streptavidin-647. After washout, neurons were subjected to electrical field stimulation (35 Hz, 0.1 s). Representative images show live-labelled APP/C99-positive puncta (**L***-top*, 4G8+, *yellow*), evoked Syp-pHluorin responses (**L***-middle*), and NH4Cl-induced fluorescence revealing the total vesicle pool (**L***-lower*); displayed using a Green Fire Blue LUT (0–255 intensity scale). Scale bars, 5 µm. **(M)** Box plots show evoked synaptic vesicle release normalized to baseline fluorescence and the total vesicle pool according to: % Evoked release = [(FL_evoked_ − FL_baseline_)/FL_NH4Cl_] × 100. Center line, median; box, interquartile range (IQR); whiskers, minimum and maximum values within 1.5× IQR; points represent N = 3 independent transfected cultures. Statistical significance was determined using one-way ANOVA with post hoc multiple-comparison testing. Comparisons among APP/C99 variants are denoted by asterisks (*), whereas comparisons relative to mock controls are denoted by hash symbols (^#^). ^ns^ *p* > 0.05, *^/#^ *p* < 0.05, **^/##^ *p* < 0.01, ***^/###^ *p* < 0.001.

γ-Secretase inhibition (GSI; 10 μM DAPT, 4 h) preferentially increased synaptic accumulation of all C99 species at pan-synaptic boutons, defined by Syn1-positive puncta (Fig. 1C-D, S1E-F). Quantification of the GSI-induced change relative to vehicle/untreated conditions revealed differential accumulation among the APP/C99 variants (Fig. 1D). C99^YENPTY^ exhibited the largest increase in synaptic accumulation following GSI treatment. Similarly, C99^ANAETA^ and C99^ΔCT^ accumulated and showed a modest increase. The lowest increase in APP-FL neurons could be attributed to GSI treatment only minimally altering APP-FL, but rather APP-CTFs levels. In our previous work, we demonstrated that APP and components of the amyloidogenic processing machinery preferentially localize to excitatory glutamatergic synapses (Kapadia *et al*, 2025). Thus, we next assessed localization specifically within excitatory vGLUT1 presynaptic boutons (Fig. 1E-F). Co-localization with vGLUT1-positive puncta showed that all APP-FL and C99 variants localized to excitatory synapses, and their overlap significantly increased upon GSI treatment (Fig. 1G and S1F). Here, C99^YENPTY^ showed the highest accumulation, followed by C99^ANAETA^, C99^ΔCT^ and APP-FL, respectively (Fig. 1G).

These observations suggest that APP-CTFβ is present within excitatory presynaptic terminals (Fig. 1B, F) and accumulate upon γ-secretase inhibition (Fig. 1D, G). The intracellular C-terminal domain may facilitate interactions with trafficking adaptors, possibly via the ‘YENPTY’ binding motif, vesicle-associated proteins, or membrane lipids that aid in stabilizing APP-CTFβ at glutamatergic presynapses. Loss of these interactions could possibly explain why C99^ANAETA^ and C99^ΔCT^ show only a modest increase (Fig. 1D, G). On the other hand, the modest increase in the case of APP-FL could be attributed to the major increase in APP-CTFα, alongside APP-CTFβ species.

Previous studies have demonstrated that accumulation of APP-CTFβ at excitatory presynaptic sites influences synaptic activity (Kapadia *et al*, 2025; Das *et al*, 2021; Tambini *et al*, 2019; Yao *et al*, 2019). To test this possibility, we examined levels of glutamate in conditioned media (Fig. 1H-J), as a proxy for neurotransmitter release. In untreated conditions, expression of C99^YENPTY^ and C99^ANAETA^ variants led to a modest, yet significant, increase in glutamate levels in conditioned media, compared to mock conditions (Fig. 1I). Here, the expression of APP-FL or C99^ΔCT^-the variant lacking the entire intracellular C-terminal domain, did not change the global glutamate levels (Fig. 1I). Upon GSI treatment, C99^YENPTY^ showed the highest change, followed by C99^ANAETA^ and APP-FL, wherein, changes for C99^ΔCT^ were insignificant, compared to mock controls (Fig. 1J). The intermediate phenotype observed for APP-FL is consistent with the generation of a heterogeneous pool of APP-derived C-terminal fragments upon γ- secretase inhibition, including both APP-CTFα and APP-CTFβ, whereas the C99 constructs selectively enrich for APP-CTFβ species. In this experiment, it is important to take into consideration the dilution effects of mixed neuronal cultures as mock cells also showed an increase in glutamate levels upon γ-secretase inhibition (Fig. 1J; *dotted line*). These observations indicate that APP-CTFβ accumulation, particularly in the presence of an intact intracellular C-terminal domain, is associated with enhanced glutamate release.

Next, to specifically isolate effects from APP/C99 containing synapses, we measured synaptic vesicle release using synaptophysin (Syp)-pHluorin imaging (Fig. 1K-M). Neurons were treated with or without GSI in the presence of tetrodotoxin (TTX) to isolate evoked and asynchronous release components. Changes in fluorescence values were normalised to the baseline and total vesicle pool defined by NH₄Cl application. Here, we computed values from ROIs of neuronal processes positive for pHlourin and live labelled 4G8+ puncta (Fig. 1L).

Among all constructs, C99^YENPTY^ produced the largest increase in evoked release, followed by APP-FL and C99^ANAETA^, which significantly increased upon GSI treatment. The deletion of the intracellular C-terminal domain abolished this effect, as C99^ΔCT^ showed little or no effect, and was comparable to mock and untreated conditions (Fig. 1M). Post-imaging, neurons were fixed, stained and imaged, confirming that surface labelled 4G8+ puncta detected for C99^YENPTY^, APP-FL, C99^ANAETA^, and least C99^ΔCT^ colocalise with presynaptic boutons stained with Syn1/2 (Fig. S1G). The stronger effect in case of C99^YENPTY^, followed by C99^ANAETA^, and a minimal effect in case of C99^ΔCT^, suggests that localisation and the functional gain of presynaptic vesicle release depend on the intracellular C-terminal domain of APP-CTFβ, but is largely independent of the YENPTY motif. Together, these observations demonstrate that APP-CTFβ accumulation at excitatory presynaptic terminals correlates with enhanced evoked synaptic vesicle release and requires an intact intracellular C-terminal domain.

### γ-secretase-dependent accumulation of APP-CTFβ promotes its oligomerization

Although the C-terminal domain was required for APP-CTFβ-mediated enhancement of synaptic vesicle release (Fig. 1), the marked increase in activity following γ-secretase inhibition suggested that APP-CTFβ accumulation alone may not fully account for this phenotype. Aβ has long been shown to undergo self-assembly into oligomeric species, which are considered major contributors to synaptic dysfunction and toxicity in AD (Viola & Klein, 2015; Sakono & Zako, 2010; Walsh & Selkoe, 2007; Sciaccaluga *et al*, 2021; Hayden & Teplow, 2013; Cavallucci *et al*, 2012; Cline *et al*, 2018). More recent studies have further demonstrated that APP-CTFβ can similarly form dimers and higher order assemblies (Pantelopulos *et al*, 2022, 2018; Badot *et al*, 2026; Perrin *et al*, 2020; Vaillant-Beuchot *et al*, 2021). In addition, previous computational modelling studies suggested that APP-CTFβ interactions with synaptic vesicles are strongly influenced by its multimerization state, i.e., dimers and trimers (Kapadia *et al*, 2025). Therefore, we next asked whether accumulation of APP-CTFβ promotes a transition from monomeric to oligomeric states.

To address this, we generated CHO cell lines expressing human APP-FL and the three C99 variants (Fig. 2A) and examined their localization and assembly under γ-secretase inhibition (GSI; 10 μM DAPT, 16-18 h) treatment conditions (Fig. 2). Immunofluorescence analysis revealed membrane and intracellular localisation of APP-FL and all C99 variants (Fig. 2B; Fig. S2E), consistent with their transmembrane nature. Notably, C99^ΔCT^ displayed enhanced intracellular retention, consistent with observed somatic accumulation in primary neurons (Fig. 1A, C; S1E).

**Figure 2.**
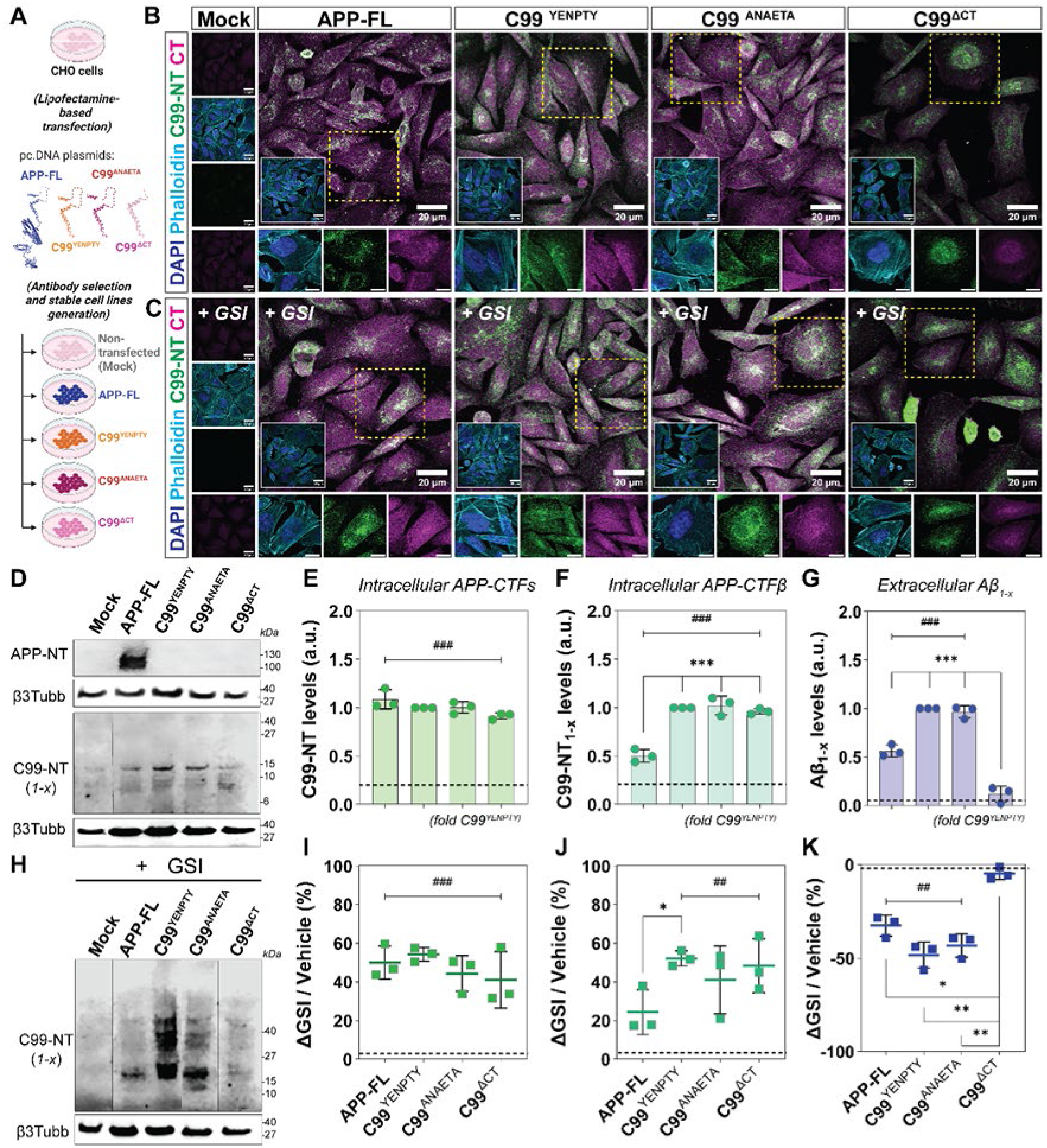
γ-Secretase inhibition promotes APP-CTFβ accumulation and higher-order assembly formation in transfected CHO cells. **(A)** Schematic of experimental workflow for transfection of CHO cells with APP-FL or C99 variants, followed by generation of stable cell lines (via G418 antibiotic selection) for downstream experimental analyses. Non-transfected condition (lipofectamine without DNA) is indicated as mock. (**B-C**) Representative immunocytochemistry images of CHO cells expressing APP-FL, C99^YENPTY^, C99^ANAETA^, or C99^ΔCT^; following vehicle (**B**) and GSI treated (+ GSI, **C**) conditions. APP/C99-species were detected using 4G8 (*green*) and co-stained with C-terminal antibody (C1/6.1, *magenta*), DAPI (*blue*), and phalloidin (*cyan*). Insets show higher magnification views. Scale bars, 20 µm; zoom-in panels, 10 µm. (**D**) Western immunoblots (WB) confirming expression of APP-FL and C99 variants in stable CHO cell lysates under basal conditions. Immunoblots depict APP N-terminal (antibody: APP-NT) and C99-NT_1-x_ species (antibody: 82E1) with β3-tubulin as respective loading control. (**E-F**) Bar plots depict quantification of intracellular APP-CTFs (**E**, antibody clone; 4G8, WB), C99 species (**F**, antibody clone: 82E1, WB) from APP-FL- and C99-expressing stable CHO cells under basal conditions. Individual points represent average from each experiment normalised to C99^YENPTY^ condition within each biological replicate; bars, mean; error bars, S.D., dotted line, mock; N = 3 independent cultures. (**G**) ELISA analysis depicting extracellularly secreted Aβ_1-x_ species (antibody clone: 82E1) in conditioned media from APP-FL- and C99-expressing stable CHO cells under basal conditions. Individual points represent average from each experiment normalised to C99^YENPTY^ condition within each biological replicate; bars, mean; error bars, S.D., dotted line, mock; N = 3 independent cultures. (**H**) Western immunoblots showing accumulation of intracellular C99-NT_1-x_ (APP-CTFβ, antibody: 82E1) species and the appearance of higher molecular weight immunoreactive bands following γ-secretase inhibition (+GSI) in APP/C99 expressing CHO cells. β3-tubulin was used as loading control. (**I-K**) Scatter blots depicting percentage change (Δ) of intracellular APP-CTFs (**I**, antibody clone; 4G8; WB), APP-CTFβ (**J**, antibody clone: 82E1; WB) and extracellular Aβ levels (**K**, antibody clone: 82E1; ELISA), following GSI treatment relative to the corresponding vehicle controls. ΔGSI/Vehicle (%) = [(Signal_GSI_ - Signal_Vehicle_) / Signal_Vehicle_] ×100. Individual points represent average from each independent experiment; bold line, mean; error bars, S.D., dotted line, mock; N = 3 independent transfected cultures. Statistical significance was determined using one-way ANOVA with post hoc multiple-comparison testing. Comparisons among APP/C99 variants are denoted by asterisks (*), whereas comparisons relative to mock controls are denoted by hash symbols (^#^). ^ns^ *p* > 0.05, *^/#^ *p* < 0.05, **^/##^ *p* < 0.01, ***^/###^ *p* < 0.001.

Biochemical characterization confirmed robust expression of APP-FL (detected *via* APP N-terminal ectodomain; Fig. S2D, ELISA) and C99 variants (detected via N-terminal 1-x epitope of C99) in CHO cells (Fig. 2D). In basal conditions, total levels of APP/C99 species are comparable across the tested conditions (Fig. 2E, 4G8 antibody), whereas levels of APP-CTFβ is comparable between C99^YENPTY^, followed by C99^ANAETA^ and C99^ΔCT^, and the least for APP-FL (Fig. 2F). Lastly, C99^YENPTY^, and C99^ANAETA^ produced comparable levels of Aβ, with lower levels for APP-FL as it is also concurrently processed through the non-amyloidogenic pathway (Aβ_1-x_: 82E1, Fig. 2G; and total Aβ: 2964, Fig. S2A). In contrast, secreted Aβ species were largely absent in C99^ΔCT^ expressing cells. Here, the absence of C-terminal domain plausibly prohibits the release of Aβ species (Fig. 2G and S2A).

As expected, γ-secretase inhibition significantly decreased extracellular Aβ species (Aβ_1-x_: 82E1, Fig. 2K; and total Aβ: 2964, Fig. S2B) for C99^YENPTY^ and C99^ANAETA^, followed by APP-FL expressing cells. C99^ΔCT^ showed minimal or no effect in the levels of secreted Aβ species as compared to mock controls. Now, GSI treatment led to a significant increase in APP-CTFβ/s species across all C99 variants and APP-FL conditions, when compared to mock conditions (Fig. 2H-I). Particularly, APP-CTFβ is significantly enriched for all C99 variants, along with a moderate increase in APP-FL expressing cells (Fig. 2J). Strikingly, immunoblot analysis revealed the appearance of higher molecular weight 82E1-positive assemblies following GSI treatment, most prominently in C99^YENPTY^ and C99^ANAETA^ expressing cells, whereas APP-FL derived CTFs and C99^ΔCT^ showed minimal or weaker signals (Fig. 2H; Fig. S3A).

To determine whether these assemblies represented oligomeric APP-CTFβ species, we probed native PAGE immunoblots using both, the 4G8 (Fig. 3A, *top*) and the oligomer conformation-specific antibody-A11 (Fig. 3A, *bottom*). In parallel, immunocytochemistry on GSI treated C99^YENPTY^ expressing cells show an increase in A11+ punctate staining pattern both on the membrane as well as intracellular compartments, respectively (Fig. 3B). C99^ANAETA^, APP-FL and C99^ΔCT^ showed modest changes in this regard (Fig. S3B). To quantitatively assess oligomer formation, we performed single-molecule pull-down (SIMPull; Fig. 3C-D, Fig. S3C) and in-house sandwich ELISA (Fig. 3E-F) assays using 4G8 as a capture antibody and detection with the oligomer-specific A11 antibody. Under basal conditions, APP-FL and C99 variants exhibited low and comparable levels of A11-positive species. In contrast, GSI treatment resulted in a pronounced increase in oligomeric assemblies for C99^YENPTY^ (5,7-fold), C99^ANAETA^ (3,2-fold), and APP-FL (1,2-fold); whereas changes in C99^ΔCT^ remains negligible (Fig. 3D and S3C). Here Aβ_42_ oligomers were used as positive normalisation controls (Fig. S3C). Consistent results were obtained using ELISA experiments, wherein A11-positive puncta were strongly enriched under GSI conditions in C99^YENPTY^ and C99^ANAETA^ expressing cells (Fig. 3F). The lower signals seen in the case of APP-FL could be because of the lower proportion of APP-CTFβ species among the total APP-CTFs pool. Since changes for C99^ΔCT^ remain negligible or quite modest, we can infer that the loss of C-terminal domain could plausibly induce changes in the transmembrane helix altering its biophysical properties and/or its localisation defects as seen earlier.

**Figure 3.**
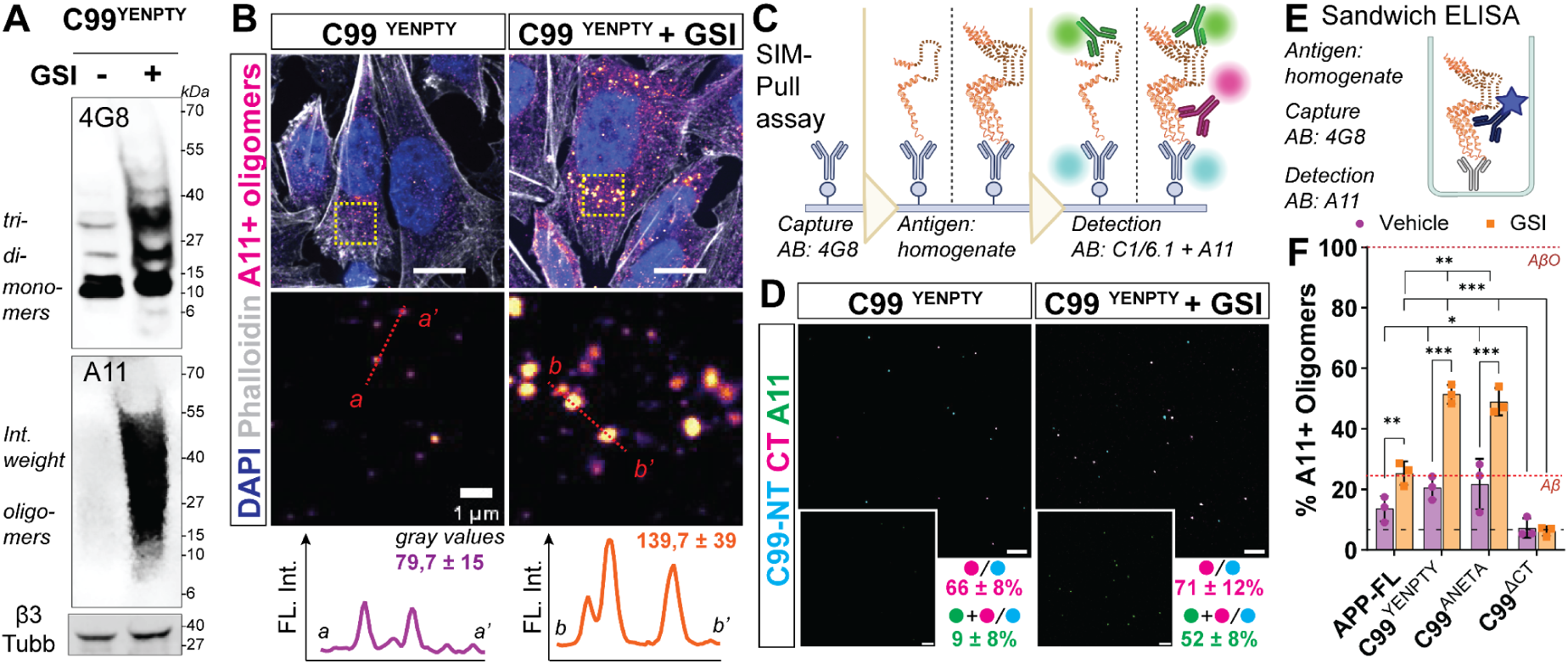
γ-Secretase inhibition promotes APP-CTFβ oligomerization detected by complementary biochemical, imaging, and single-molecule approaches. **(A)** Native western immunoblots of CHO cells expressing C99^YENPTY^ under untreated (vehicle, -GSI) and treated (+GSI) conditions. Detection with 4G8 reveals monomeric, dimeric, trimeric, and higher molecular weight APP-CTFβ species that accumulate following γ-secretase inhibition. The conformation-specific A11 antibody detects oligomeric assemblies that become prominent following γ-secretase inhibition. β3-tubulin served as loading control. **(B)** Representative immunocytochemistry images of CHO cells expressing C99^YENPTY^ under GSI treated and untreated conditions, stained with the oligomer-specific A11 antibody. Graph insets show representative fluorescence intensity profiles across A11-positive puncta. Scale bars, 10 µm; zoom-in panels, 1 µm **(C)** Schematic of the single-molecule pull-down (SIMPull) assay. APP/C99 species present in CHO cell homogenates act as antigens and were captured using 4G8 antibody (*blue*). C-terminal epitopes were detected with C1/6.1 (*magenta*) and the oligomers with conformation-specific A11 antibody (*green*). **(D)** Representative SIMPull images showing A11-positive puncta (*green*) detected from homogenates C99^YENPTY^-expressing cells under vehicle and +GSI conditions. Scale bars, 5 µm. Numerical values indicate the percentage of 4G8-captured puncta that were positive for C1/6.1 alone (*magenta*) or simultaneously positive for C1/6.1 and A11 (*green*). **(E)** Schematic representation of the oligomer-specific sandwich ELISA, using 4G8 as capture, GSI treated CHO cell homogenate as antigen, and A11 as detection antibody. **(F)** Bar plots depict quantification of A11-positive ELISA signals across APP-FL and C99 variants under vehicle (*magenta*) and GSI treatment (*orange*) conditions. A11-reactive Aβ_42_ oligomers (AβO) were used as a positive control and assigned a value of 100% (*maroon dotted line*); monomeric Aβ_42_, *red dotted line*. Individual points represent technical replicates (n = 6); bars represent mean ± S.D.; N = 3 independent cultures; dotted line, mock-vehicle, bold line, mock-GSI. Statistical significance was determined using one-way ANOVA with post hoc multiple-comparison testing. Comparisons among APP/C99 variants are denoted by asterisks (*), whereas comparisons relative to mock controls are denoted by hash symbols (^#^). ^ns^ *p* > 0.05, *^/#^ *p* < 0.05, **^/##^ *p* < 0.01, ***^/###^ *p* < 0.001.

Together, these findings demonstrate that γ-secretase-dependent accumulation of APP-CTFβ promotes its assembly into oligomeric membrane-associated species, independently of canonical YENPTY-mediated interactions.

### APP-CTFβ oligomerization is associated with changes in membrane lipid organization

Given that APP-CTFβ oligomers localize to membranes, could its oligomerization state alter the lipid organization in its vicinity. Indeed, APP-FL is known to interact with lipids such as cholesterols (Wang *et al*, 2019; Beel *et al*, 2008; Marzolo & Bu, 2009; Fabelo *et al*, 2014; Hicks *et al*, 2012). Could the N-terminal ectodomain and C-terminal domain differentially influence the positioning of the transmembrane helix, thus altering local lipid environment? To test this, we generated giant plasma membrane-derived vesicles (GPMVs) from CHO cells expressing APP-FL and C99 variants (Fig. 4A-B, S4A-B). Fluorescence imaging confirmed the vesicle membrane stained with lipid markers, including BODIPY, filipin, and WGA glycosylated membrane proteins (Fig. 4C, S4B), as well as the presence of APP/C99 species, stained with 4G8 antibody (Fig. 4C). Quantitative analysis using AmplexRed cholesterol assays revealed that cholesterol levels were significantly altered in all APP/C99 cell-derived GPMVs as compared to mock controls. While the levels of cholesterol in GPMVs were lower in all C99 variants, APP-FL GPMVs showed a significant increase as compared to other conditions (Fig. 4D and S4C). In parallel, APP/C99 levels displayed differential partitioning between whole-cell lysates and isolated GPMVs (Fig. 4E, S4D). APP-FL was relatively enriched in GPMVs, whereas C99^YENPTY^ and C99^ANAETA^ were reduced compared with their respective cellular levels. C99^ΔCT^ exhibited only minor changes. These findings suggest that the extracellular and intracellular domains contribute to the membrane partitioning behaviour of APP-derived species and indicate that APP-FL and APP-CTFβ occupy distinct membrane environments. Because GPMVs preserve native membrane composition while excluding most intracellular organelles, these observations further support the notion that APP-FL and APP-CTFβ differentially associate with specific lipid microdomains. These findings are consistent with previous reports indicating the localisation of APP in cholesterol-rich domains.

**Figure 4.**
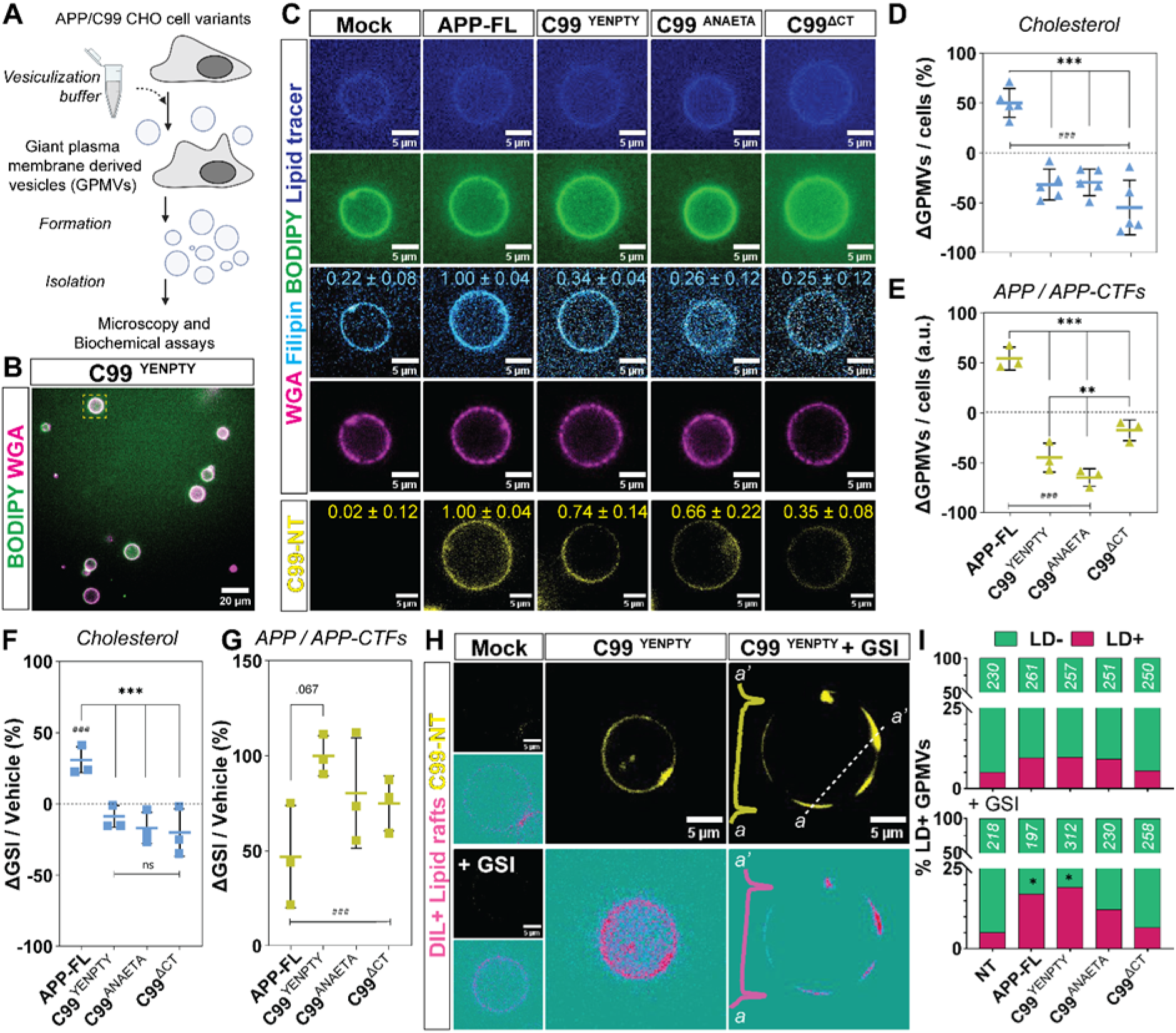
APP-CTFβ oligomerization is associated with alterations in membrane lipid organization. **(A)** Schematic of giant plasma membrane vesicle (GPMV) preparation from APP-FL- and C99-expressing CHO cells for membrane lipid analyses. **(B)** Representative images of GPMVs labeled BODIPY (lipids, *green*) and Alexa-546 labelled WGA (glycosylated membrane proteins, *magenta*) illustrating membrane integrity and vesicle morphology. Scale bars, 20 µm. **(C)** Representative fluorescence images of GPMVs derived from APP-FL- and C99-expressing CHO cells labelled with lipid tracer (*blue*), BODIPY (*green*), filipin probe (cholesterol, *cyan*), WGA (*magenta*), and APP/C99 species detected with 4G8 (*yellow*), respectively. Scale bars, 5 µm. Numerical values indicate relative fluorescence intensities normalized to APP-FL-derived GPMVs, displayed within each image, respectively. (**D-E**) Relative enrichment or depletion of cholesterol in GPMVs compared with corresponding whole-cell lysates measured using the Amplex Red cholesterol assay. Similarly, relative enrichment or depletion of APP and APP-CTF species in GPMVs compared with corresponding whole-cell lysates measured by ELISA. Scatter blots depicting percentage change (Δ) of cholesterol (**D**) and APP-CTFs (**E**) in GPMVs and lysates, respectively. ΔGPMVs/cells (%) = [(Signal_GPMVs_ - Signal_Cells_) / Signal_Cells_] ×100. Individual points represent average from each independent experiment, normalized to the corresponding mock condition, respectively. Bold line, mean; N = 3 independent cultures. (**F-G**) GPMVs were prepared from APP/C99-expressing cells following vehicle or GSI pretreatment. Relative changes in cholesterol (F) and APP/APP-CTF levels (G) in GPMVs were quantified and expressed relative to the corresponding vehicle controls. Scatter blots depicting % change (Δ) of cholesterol (**F**) and APP-CTFs (**G**) in GPMVs from APP/C99 expressing cells in untreated and GSI pretreated cells, respectively; computed as ΔGSI/Vehicle (%) = [(Values_GPMVs-GSI_ – Values_GPMVs-Vehicle_) / Values_GPMVs-Vehicle_] ×100. Individual points represent average from each independent experiment. Bold line, mean; N = 3 independent cultures. **(H)** Representative GPMVs labeled with APP/C99 (4G8) and lipid domain marker (DiL) under vehicle and GSI treated (+ GSI) conditions. N = 2 independent experiments. Scale bars, 5 µm. Fluorescence intensity profiles (a-a′) illustrate redistribution of APP/C99 species relative to membrane lipid domains. **(I)** Quantification of GPMVs displaying low (LD−) or high (LD+) lipid-disordered membrane domains across APP/C99 variants under vehicle and GSI-treated conditions. Numbers within bars indicate total GPMVs analyzed, respectively. N = 2 independent experiments. Statistical significance was determined using one-way ANOVA with post hoc multiple-comparison testing. Comparisons among APP/C99 variants are denoted by asterisks (*), whereas comparisons relative to mock controls are denoted by hash symbols (^#^). ^ns^ *p* > 0.05, *^/#^ *p* < 0.05, **^/##^ *p* < 0.01, ***^/###^ *p* < 0.001.

Next, we compared GPMVs that were derived from cells presubjected to γ-secretase inhibition to that of untreated/vehicle conditions. Cholesterol levels were augmented in the case of APP-FL conditions, but in the case of all C99-cells derived GPMVs, they remained quite comparable, if only slightly decreased as compared to non-transfected controls (Fig. 4F, S5A). The relative changes in the levels of 4G8 positive species within the GPMVs (Fig. 4G, S5B), showed a similar increase as seen in the case of the cells (Fig. 2E, I). Here, while APP/C99 levels in vehicle conditions remained comparable, they significantly increased in all conditions with highest changes in C99^YENPTY^, followed by C99^ANAETA^, C99^ΔCT^ and least in APP-FL (Fig. 4G). Together, these results suggest that while GSI treatment robustly increases APP-CTF accumulation, changes in cholesterol partitioning are largely driven by APP-FL and are not proportional to APP-CTF abundance.

Following γ-secretase inhibition, C99^YENPTY^ expressing cells exhibited significant alterations in the distribution of lipid domains, as assessed by co-staining with lipid ordered-phase domain (LD) markers (Fig. 4H-I, S5C-D). These changes were most prominent in C99^YENPTY^ conditions, APP-FL and C99^ANAETA^ displayed similar but lower probabilities of this occurrence, whereas C99^ΔCT^ showed little or no effect (Fig. 4H-I, S5C-D). Here, imaging-based analyses demonstrated pronounced changes in lipid domain organization, whereas biochemical analyses revealed only modest alterations in total cholesterol content across most C99 conditions (Fig. 4F-G, S4C-D). The emergence of ordered lipid domains following GSI treatment is dependent on the abundance of C99 species, particularly C99^YENPTY^, despite relatively modest changes in total cholesterol content. This emergence of ordered membrane domains cannot be explained solely by changes in bulk cholesterol abundance but rather reflects a C99-dependent redistribution of membrane lipids into spatially distinct microdomains.

In APP-FL-expressing cells, increases in both cholesterol and APP-derived species likely reflect the accumulation of multiple APP processing intermediates following γ-secretase inhibition. Notably, C99^ΔCT^ exhibited little or no effect on membrane domain organization, consistent with its markedly reduced oligomerization propensity (Fig. 3 and S3). Here, the deletion of the C-terminal domain may alter the structural organization of the transmembrane helix and consequently impair C99^ΔCT^-mediated interactions with surrounding membrane lipids, although the precise mechanism remains to be determined. In contrast, C99^YENPTY^ and C99^ANAETA^ exhibited substantial membrane reorganization despite comparatively minor changes in total cholesterol levels, suggesting that APP-CTFβ accumulation and oligomerization are sufficient to alter local membrane organization. Together, these findings indicate that γ-secretase-dependent APP-CTFβ oligomerization is accompanied by a reorganization of membrane lipids and promotes the formation of ordered membrane domains.

### APP-CTFβ oligomers promote synaptic vesicle tethering via the C-terminal domain

Having shown that γ-secretase-dependent APP-CTFβ accumulation promotes oligomerization and membrane reorganization, we next asked whether these assemblies directly influence synaptic vesicle recruitment. To test this, we used an extended SIMPull-based vesicle tethering assay, in which APP/C99 species were captured on 4G8 antibody-functionalized surfaces and incubated with fluorescently labelled synaptic vesicles (Fig. 5A, B).

**Figure 5.**
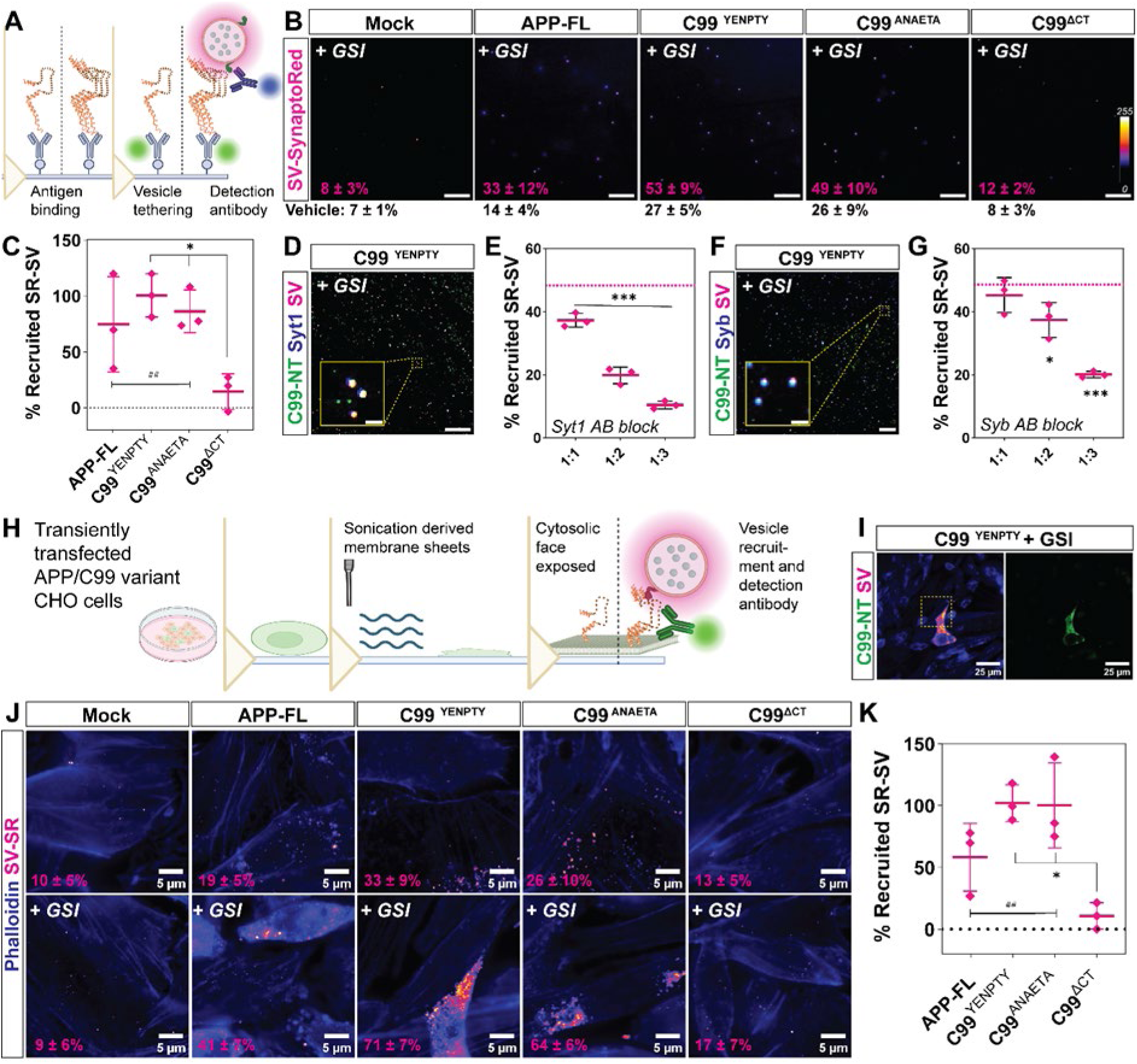
APP-CTFβ oligomers enhance synaptic vesicle tethering via the C-terminal domain. **(A)** Schematic of the extended SIMPull -based synaptic vesicle tethering assay. APP/C99-derived species are captured on antibody-functionalized glass surfaces, followed by incubation with fluorescently labelled synaptic vesicles with SynaptoRed (SV-SR, *magenta*), allowing direct quantification of vesicle recruitment. **(B)** Representative SIMPull images showing recruitment of SynaptoRed-labelled synaptic vesicles (SV-SR, displayed using a Fire LUT) to APP/C99-derived species captured from CHO cell homogenates under vehicle and γ-secretase inhibited (+GSI) conditions. Scale bars, 10 µm. Numerical values indicate the percentage of recruited SV-SR puncta (mean ± S.D.) under vehicle (*black*) and GSI-treated (*magenta*) conditions; N = 3 independent experiments. **(C)** Quantification of GSI-dependent synaptic vesicle recruitment measured by SIMPull. For each construct, vesicle recruitment under vehicle conditions was subtracted from the corresponding GSI-treated condition (ΔGSI = GSI − Vehicle). Values were subsequently normalized to the mean response obtained for C99^YENPTY^, which was set to 100%. Individual points represent independent experiments; bold line, mean ± S.D.; N = 3 independent experiments. (**D, F**) Representative SIMPull images demonstrating colocalization of tethered synaptic vesicles with synaptotagmin-1 (Syt1, **D**) and synaptobrevins (Syb, **F**), confirming recruitment of intact synaptic vesicles containing canonical vesicle-associated proteins. Insets show higher magnification views. Scale bars, 10 µm; zoomed-in panels, 1 µm. (**E, G**) Quantification of synaptic vesicle recruitment following antibody-mediated blockade of Syt1 (E) or Syb (G). Progressive reduction in vesicle tethering upon increasing antibody concentrations (1:1, 1:2 and 1:3 dilution ratios) indicates that APP-CTFβ-mediated vesicle recruitment depends on C-terminal interactions with vesicle-associated SNARE machinery. Dotted lines indicate recruitment levels in the absence of blocking antibodies. Individual points represent independent experiments; bold line, mean ± S.D.; Bold line, mean; N = 3 independent experiments. **(H)** Schematic of the membrane sheet-based tethering assay. Sonication-derived plasma membrane sheets expose the cytosolic membrane surface, enabling assessment of APP/C99-mediated synaptic vesicle recruitment in a membrane-associated environment. **(I)** Representative membrane sheet derived from C99^YENPTY^-expressing CHO cells following GSI treatment. C99 species were detected with 4G8 antibody (*green*), and recruited synaptic vesicles were loaded with SynaptoRed (*magenta*). Scale bars, 25 µm. **(J)** Representative membrane sheet tethering assay images from mock, APP-FL, and C99-expressing CHO cells under vehicle and GSI-treated (+GSI) conditions. Recruited SynaptoRed-labelled synaptic vesicles are displayed using a Fire LUT; membrane sheets are visualized with phalloidin (*blue*). Scale bars, 5 µm. **(K)** Quantification of GSI-dependent synaptic vesicle recruitment in membrane sheet assays. For each construct, recruitment measured under vehicle conditions was subtracted from the corresponding GSI-treated condition (ΔGSI = GSI − Vehicle). Values were subsequently normalized to the mean response obtained for C99^YENPTY^, which was set to 100%. Individual points represent independent experiments; bold line, mean ± S.D.; N = 3 independent experiments. Statistical significance was determined using one-way ANOVA with post hoc multiple-comparison testing. Comparisons among APP/C99 variants are denoted by asterisks (*), whereas comparisons relative to mock controls are denoted by hash symbols (^#^). ^ns^ *p* > 0.05, *^/#^ *p* < 0.05, **^/##^ *p* < 0.01, ***^/###^ *p* < 0.001.

Under basal conditions, vesicle tethering was low across most conditions, with only modest recruitment observed for C99^YENPTY^ and C99^ANAETA^ (Fig. 5B, *values in vehicle conditions*). Following γ-secretase inhibition, C99^YENPTY^ and C99^ANAETA^ showed a pronounced increase in synaptic vesicle tethering, whereas APP-FL displayed a weaker effect and C99^ΔCT^ remained largely inactive (Fig. 5B, C). This pattern closely mirrored the oligomerization profile (Fig. 3) and evoked release measurements (Fig. 1), suggesting that APP-CTFβ oligomeric assemblies are associated with synaptic activity by enhancing vesicle recruitment.

To confirm that the recruited puncta represented intact synaptic vesicles, we immunolabelled tethered vesicles for synaptotagmin-1 (Syt1) and synaptobrevin/VAMPs (Syb). Recruited vesicles strongly colocalized with both markers, indicating specific capture of synaptic vesicle particles (Fig. 5D, F). As shown previously, APP-CTFβ interacts with these synaptic vesicle proteins via its C-terminus. Here, antibody-mediated blocking of Syt1 or Syb epitopes reduced vesicle tethering in an antibody dose-dependent manner (Fig. 5E, G), supporting the involvement of canonical synaptic vesicle proteins in APP-CTFβ-mediated recruitment.

To determine whether this interaction also occurs in neurons, we performed SIMPull-based vesicle tethering assays using homogenates from GSI-treated neuronal cultures expressing APP-FL and C99^YENPTY^ (Fig. S6A-B). C99^YENPTY^ expressing neurons showed an approx. ∼2-fold enrichment in tethered SR-SV puncta upon GSI treatment (Fig. S6A-B), supporting the relevance of this mechanism in a neuronal context. Here, preformed Aβ_42_ oligomers showed no effect, comparable to that of C99^ΔCT^ conditions. These findings confirm the recruitment of synaptic vesicles is dependent on the C-terminal domain, wherein the specific amino acid(s) or sequences mediating these interactions remains to be investigated.

Finally, we validated these findings in a membrane context, we performed vesicle tethering assays on sonication-derived membrane sheets (Perez et al., 2006), selectively exposing the cytosolic face (Fig. 5H-I, S6C). The use of transiently transfected cells (Fig. S2C, E) allowed vesicle recruitment to be compared directly between transfected and non-transfected cells within the same image, providing an in-image control (Fig. 5I, S6D). Consistent with SIMPull results, membrane sheets derived from C99^YENPTY^ and C99^ANAETA^ expressing cells showed vesicle recruitment, whereas APP-FL displayed a weaker effect and C99^ΔCT^ remained largely inactive (Fig. 5J). Upon γ-secretase inhibition, this effect was significantly augmented (Fig. 5I-K). Importantly, the comparable behaviour of C99^YENPTY^ and C99^ANAETA^ indicates that vesicle tethering is independent of the YENPTY motif, but strictly requires the C-terminal domain, as deletion of this region abolishes the interaction. Together, these results demonstrate that APP-CTFβ oligomers act as membrane-associated scaffolds that promote synaptic vesicle tethering *via* C-terminal domain-dependent interactions with synaptic vesicle proteins.

## Discussion

In this study, we identify APP-CTFβ as a presynaptic gain-of-function regulator that alters synaptic function. We show that accumulation of APP-CTFβ promotes its self-assembly and oligomerization, leading to membrane remodelling and synaptic vesicle tethering *via* its C-terminal domain. Interestingly, APP-FL exhibited only modest effects across all assays, whereas C99^ANAETA^ largely phenocopied C99^YENPTY^ despite disruption of the canonical binding motif. These observations indicate that generation, accumulation, and oligomerisation of APP-CTFβ, rather than full-length APP itself, is a critical determinant of this gain-of-function properties. Additionally, APP-CTFβ oligomerization and vesicle tethering are largely independent of YENPTY-mediated adaptor interactions. These findings challenge the traditional view of APP-CTFβ as a transient intermediate/by-product but instead identify it as a signalling entity modulating presynaptic function, with a potential implication in the hyperexcitability phase in early AD onset (Anastacio *et al*, 2022; Das *et al*, 2021; Kapadia *et al*, 2025; Tambini *et al*, 2019; Yao *et al*, 2019).

In neurons, acute γ-secretase inhibition increased presynaptic activity by enhancing vesicle release, whereas deletion of the C-terminal domain abolished this effect. Importantly, the gain of presynaptic function could be attributed to an increase in the local concentration of APP-CTFβ species upon γ-secretase inhibition. Mechanistically, we demonstrate that accumulation of APP-CTFβ facilitates the formation of higher-order, A11-reactive oligomers. These assemblies are strongly associated with enhanced synaptic vesicle tethering in both single-molecule and membrane-based assays. We can hypothesize that monomeric APP-CTFβ has limited vesicle-binding capacity, whereas oligomerization likely increases the local valency and avidity of its C-terminal interaction. These multimeric platforms could simultaneously engage with multiple vesicle-associated proteins. Such a mechanism could explain why vesicle tethering increases disproportionately following γ-secretase inhibition, locally increasing vesicle density and facilitating release.

Notably, the A11 reactivity of APP-CTFβ oligomers suggests structural similarities to prefibrillar oligomeric assemblies previously described for Aβ, which have been widely implicated in synaptic dysfunction and neuronal toxicity (Cai *et al*, 2023; Cerpa *et al*, 2008; Giorgio *et al*, 2024; Martinsson *et al*, 2022; Rijal Upadhaya *et al*, 2012; Sciaccaluga *et al*, 2021; Siddu *et al*, 2025; Takahashi *et al*, 2004; Wei *et al*, 2010; Wilson *et al*, 2025; Zott *et al*, 2019). However, in contrast to Aβ oligomers, which could act both intracellularly and extracellularly (Cuello, 2005; Ding *et al*, 2019; Fontana *et al*, 2020; LaFerla *et al*, 2007; Oddo *et al*, 2006), APP-CTFβ oligomers being primarily membrane bound, operate intracellularly at the synaptic and neuronal membrane (Barthet *et al*, 2018; Burrinha *et al*, 2021; Daly *et al*, 1998; Essayan-Perez & Südhof, 2023; Ferrer-Raventós *et al*, 2023; Jordà-Siquier *et al*, 2022; Kapadia *et al*, 2025; Kwart *et al*, 2019; Luo *et al*, 2024; Martinsson *et al*, 2022; Schilling *et al*, 2023; Tambini *et al*, 2019; Yao *et al*, 2019; Zhou *et al*, 2022); as well as impair intracellular organelle homeostasis (Bécot, 2019; Bourgeois *et al*, 2018; Hung & Livesey, 2018; Kwart *et al*, 2019; Lauritzen *et al*, 2016, 2019b, 2012; Vaillant-Beuchot *et al*, 2021; Lauritzen *et al*, 2019a; Badot *et al*, 2026). This spatial distinction points to a fundamentally distinct mode of action, whereby APP-CTFβ oligomers could directly modulate vesicle organization and release through membrane-associated interactions. These findings suggest that oligomerization represents a shared structural feature of APP-derived membranous fragments, but with distinct functional consequences depending on their localization and context within the neuron.

APP has previously been reported to associate with cholesterol-rich membrane microdomains, often referred to as lipid rafts (Cheng *et al*, 2007; Cordy *et al*, 2006; Fabelo *et al*, 2014; Hicks *et al*, 2012; Rappoport, 2025; Rushworth & Hooper, 2010). These represent liquid-ordered (L_o_) phases enriched in cholesterol and sphingolipids, distinct from the more fluid liquid-disordered (L_d_) regions of the membrane (Cordy *et al*, 2006; Simons & Ehehalt, 9 2002; Vetrivel & Thinakaran, 2010). Such lipid domains have been implicated in the spatial organization of synaptic proteins and vesicle fusion machinery (Lingwood & Simons, 2010; Prescott *et al*, 2009; Sapoń *et al*, 2023). Particularly, APP-CTFβ oligomerization was associated with changes in local membrane lipid (re)organization.

Presynaptic vesicle docking and fusion occur within highly specialized membrane environments enriched in cholesterol, phosphoinositides and SNARE-associated proteins. In the case of APP-CTFβ oligomers, altered lipid distribution and domain formation are consistent with a redistribution rather than a global change in cholesterol levels. Consequently, local APP-CTFβ-driven membrane reorganization could indirectly facilitate vesicle recruitment by concentrating components of the release machinery within confined membrane regions. These changes could, in turn, influence vesicle recruitment by modulating membrane properties, such as its thickness or curvature, protein clustering, or the availability of key phospholipids such as PIP2 (Kapadia *et al*, 2025), although further work will be required to define the precise biophysical mechanisms underlying these effects.

Mutations in *APP* and presenilin (*PSEN1* and *PSEN2*) genes, which underlie familial early-onset AD, γ-secretase processivity, thereby increasing the relative abundance and lifetime of membrane-retained APP-CTF species (Kwart *et al*, 2019). These have been linked to neuronal dysfunction but also dysregulated synaptic activity (Annaert *et al*, 2000; Barthet *et al*, 2018; Chang & Suh, 2005; Das *et al*, 2021; Kabir *et al*, 2020; Lauritzen *et al*, 2019b; Ray *et al*, 2024; Tambini *et al*, 2019; Yao *et al*, 2019; Thordardottir *et al*, 2017). Notably, such alterations are associated with increased neuronal excitability and network hyperactivity prior to the onset of overt neurodegeneration (Giorgio *et al*, 2024; Kapadia *et al*, 2025; Lauritzen *et al*, 2019b; Martinsson *et al*, 2022). Our findings raise the possibility that accumulation and oligomerization of APP-CTFβ may represent a convergent downstream consequence of multiple genetic forms of AD.

Our findings point to a C-terminal structure-driven function of APP-CTFβ at the presynaptic membrane. These observations add in to the present mechanism from previously described roles of APP in endocytosis and intracellular signalling that implicate the YENPTY motif to mediate canonical adaptor-mediated interactions. The YENPTY motif is a well-established clathrin-mediated endocytic sorting signal (Aow *et al*, 2023; Guénette *et al*, 2017; King & Scott Turner, 2004; Nhan *et al*, 2015; Ono *et al*, 1997; Zheng & Koo, 2006), via interactions with cytosolic adaptor proteins regulating internalization of APP or its proteolytic fragments to and from the plasma membrane into the endosomal network. When the C-terminus is either deleted (C99^ΔCT^) or a mutated (C99^ANAETA^) this endocytic sorting mechanism is plausibly disrupted and fail to reach the intracellular compartments where γ-secretase cleavage is predominantly active. This is consistent with previous literature demonstrating that YENPTY-mutants fail to efficiently internalize into Rab5-positive early endosomes (Xu *et al*, 2018). This could explain the modest changes consistently observed in comparison to the C99^YENPTY^. While the contribution of the endo-lysosomal sorting machinery has not been specifically studied in this project, it would be interesting to further elucidate these mechanisms in the future.

In our experiments, we note that vesicle tethering is mediated by structural features of the intracellular C-terminal domain, with a minimal impact of the YENPTY motif. Previous studies have shown the interaction of the intracellular C-terminal domain with synaptic vesicle proteins (Del Prete *et al*, 2014; Kapadia *et al*, 2025; Kohli *et al*, 2012; Raychaudhuri & Mukhopadhyay, 2007). This interaction is much stronger for APP-CTFβ, rather than APP. The involvement of synaptotagmin and synaptobrevin suggests that APP-CTFβ oligomers can engage with core synaptic vesicle components (Barthet *et al*, 2018; Kapadia *et al*, 2025). While our data do not particularly imply direct and indirect interactions with the SNARE machinery, the reduction in vesicle tethering upon antibody-mediated blocking supports a functional dependence on these vesicle proteins. In this context, APP-CTFβ oligomers may act as membrane-associated scaffolds that increase the local availability of synaptic vesicles, thereby enhancing release probability, as observed in our experiments. These mechanisms contribute to an added role of APP-CTFβ to early network hyperexcitability observed during disease onset.

APP-FL exhibited modest effects in several assays, these were consistently weaker than those observed for APP-CTFβ variants. These findings do not exclude physiological functions of APP at synapses, rather support the idea that APP-FL primarily serves as a precursor for functionally active C-terminal fragments produced via the β-cleavage (Daly *et al*, 1998; Das *et al*, 2021; Tambini *et al*, 2019; Yao *et al*, 2019). The presence of the large extracellular component in APP-FL could plausibly induce an ‘ectodomain drag’ restricting functional signalling events by the C-terminal domain. Indeed, it has been shown that APP extracellular ectodomain regulates its proteolytic processing (Koch *et al*, 2023; Lichtenthaler, 2006; Zhang *et al*, 2021). The comparatively modest effects observed for APP-FL suggest that the large extracellular ectodomain may impose conformational or steric constraints that limit access of the C-terminal region to membrane lipids and interaction partners. Proteolytic removal of the ectodomain during β-cleavage may therefore unmask APP-CTFβ, enabling interactions with membrane-associated factors such as PIP2 and promoting its oligomerization-dependent presynaptic functions. On the other hand, the strong loss-of-function phenotype of C99^ΔCT^ further highlights the importance of the intracellular C-terminal domain for both proper localization (Kouchi *et al*, 1998) and vesicle tethering, suggesting that APP-CTFβ function depends on coordinated effects of the N-terminal epitope, the transmembrane-dependent membrane targeting and C-terminal protein interaction capacity.

While this study provides a mechanistic framework linking APP-CTFβ oligomerization to synaptic vesicle tethering, several limitations should be considered. First, although our oversimplified single-molecule and membrane reconstitution approaches enable controlled interrogation of APP-CTFβ or its oligomer-dependent vesicle interactions, they do not fully recapitulate the ultrastructural complexity present in native synapses. Future studies employing high-resolution electron microscopy or in situ approaches will be important to directly visualize vesicle organization at presynaptic boutons and the different synapse subtypes. A second limitation is that most mechanistic experiments relied on overexpression systems. Although APP-CTFβ accumulation also occurs in disease-relevant contexts and endogenous APP-CTFβ has been detected in human AD brain tissue, future studies will be required to determine the extent to which endogenous APP-CTFβ oligomerization contributes to synaptic dysfunction in vivo. Third, while our data support a functional role for Syt1 and Syb in APP-CTFβ-mediated vesicle tethering, the precise molecular interfaces and whether these interactions are direct, remain to be defined. Fourth, GPMVs represent a highly simplified model system that lacks essential cytosolic components and cytoskeletal scaffolds. Additionally, size-variability may alter spatial packing of proteins, thus the contributions partly driven by vesicle curvature variations should also be taken into consideration. Finally, our work focuses on β-cleavage-derived APP-CTFβ (C99), and it will be important to determine whether alternative APP fragments, such as the α-cleavage-derived CTFα (C83, loss of 16 amino acids from the N-terminal epitope), exhibit similar or distinct effects on oligomerisation capabilities, vesicle tethering and membrane lipid organization. Addressing these questions will further refine our understanding of how APP processing pathways differentially regulate synaptic function.

Taken together, our findings identify APP-CTFβ oligomerization as a molecular switch that couples altered APP processing to presynaptic dysfunction. Taken together, our findings support a model in which APP-CTFβ accumulation drives the formation of membrane-associated oligomeric assemblies that reorganize local membrane environments, recruit synaptic vesicles, and increase neurotransmitter release probability. This pathway provides a mechanistic framework linking aberrant APP processing to the early neuronal hyperactivity that precedes overt neurodegeneration in AD pathogenesis.

## Material and Methods

### DNA constructs

Human APP695 (APP-FL) and APP-CTFβ (C99) pc.DNA constructs used for all experiments, were obtained from Addgene (RRID: #30137, #30146, #30144, and #30143). C99 variants included wild-type C99^YENPTY^, a mutant in the YENPTY motif (C99^ANAETA^), and a C-terminal deletion mutant (C99^ΔCT^). All constructs obtained were cloned into mammalian expression vectors under a pCAX promoter to promote membrane expression. Syp-pHluorin construct (Addgene plasmid #168510; RRID: #168510).

### Neuronal and CHO cell culture

Animals were handled and maintained according to the guidelines laid down by the Animal Welfare Body (AWB) (Instantie voor Dierenwelzijn IvD) in line with the animal experimentation policy within Radboud University and RadboudUMC; under the license/protocol numbers 2021-0040-001(-004) to Dr. Anne-Sophie Hafner. Primary cortical neurons were prepared and maintained as previously described (Kapadia et al., 2025). All experiments complied with national animal care guidelines and the guidelines issued by the Radboud UMC, Nijmegen and were approved by local authorities. Briefly, cortices dissected from embryonic (day 18) rat pups of either sex (Sprague-Dawley strain; Janvier), were trypsinised (1x, 15-20 min), and mechanically dissociated in 1% normal horse serum (NHS; ThermoFischer Scientific, #16050130) containing Basal Media Eagle (Life technologies, #21010046), containing supplements like B27 plus (Gibco, #A3582801), penicillin-streptomycin (1x PS, Gibco, #15140122), non-essential amino acids (1x NEAA, ThermoFischer, #11140050), sodium pyruvate (1x, NaPy, ThermoFischer, #11360070). Dissociated cells were plated on poly-D-lysine coated (Sigma Aldrich, #A-003-E) glass coverslips (VWR, #631-1577/ 79/ 84). Neurons were maintained and matured in a humidified atmosphere at 37°C and 5% CO2 in growth medium (described above, without serum). Neurons were fed on DIV1, DIV4, DIV7, DIV14 by replacing 1:1 media replacement with BrainPhys™ neuronal medium (StemCell technologies, # 05790) supplemented with NeuroCult™ SM1 neuronal supplement (StemCell technologies, # 05711) and PS (1x). Experiments were performed during IV18-22 to ensure mature synapses and network formation. On the other hand, CHO cells (ATCC, #CCL-61) were maintained in Ham’s F-12 nutrient mix (Gibco, #11765047) supplemented with 10% fetal bovine serum (ThermoFischer, #A5209501) and penicillin-streptomycin (ThermoFischer, # 15140122).

### Transfections

Neuron transfections were performed at DIV7 using a Lipofectamine 2000 (ThermoFischer, #11668027) based protocol as previously described (Kapadia *et al*, 2025). For live imaging experiments, neurons were co-transfected with Synaptophysin-pHluorin constructs. Neurons were cultured until DIV18-22 and used for all downstream experiments. CHO cells were transiently transfected with APP-FL or C99 constructs using Lipofectamine, according to manufacturer’s protocol (ThermoFischer #11668027). Stable CHO cell lines were generated by G418 antibiotic selection (Invivogen #ant-gn-1) and maintained under selective conditions.

### Pharmacological treatments

Primary cortical neurons (DIV21–22) were treated with DAPT (10 μM; γ-secretase inhibitor, GSI; Cayman Chemicals, #13197-5), for 4 h (primary neurons) or overnight/18 h (CHO cells) in culture media. To inhibit evoked and asynchronous release, neurons were treated with TTX (HelloBio, # HB1035) (1 µM, 4 h) alone or simultaneously with GSI.

### Immunocytochemistry

Following treatment, neurons were either fixed with 4% PFA (ThermoFischer, # A11313.36) + 4% sucrose (Sigma, #S9378-1KG) in PBS (VWR, #A0965.9010) for 15 min at room temperature (RT). Next, neurons were permeabilized using permeabilization buffer (0.025% Triton X-100 (Merk, #T8787-250ML) in blocking buffer) for 2 min, followed by blocking (2.5% normal horse serum (Gibco, # #26050088), 2.5% bovine serum albumin (CarlRoth, #8076,4), 0.0125% Triton X-100 in PBS) for 1 h at RT. An antigen retrieval step was performed to enhance APP antibody binding. Primary antibodies (Table S2) were applied overnight at 4°C. Coverslips were washed three times with PBS and incubated with appropriate secondary antibodies for 2 h at RT. After washing, coverslips were mounted using Moviol mounting reagent (41.67% Glycerol, 16.67% Moviol 4-88 (CarlRoth# 0713.1) in ddH₂O).

### Synaptophysin-pHluorin imaging

Experimental protocol has been previous described (Kapadia *et al*, 2025). Neurons were transfected with Syp-pHluorin at DIV7. At DIV21, neurons were treated with GSI in the presence or absence of TTX (1 µM, 4 h), which was washed out prior to imaging. Imaging was performed at for 2,5 min, beginning with a 30 sec baseline period followed by field stimulation (30 Hz trains, 2 repetitions, 60 s intervals). To normalize for expression levels, neurons were treated with NH₄Cl solution (50 mM NH₄Cl replacing an equimolar amount of NaCl in Tyrode) for 5 min to alkalinize acidic compartments and reveal total pHluorin fluorescence (F_NH₄Cl). Lastly, neurons were fixed and processed for immunocytochemistry.

### Glutamate release assay

Extracellular glutamate levels were measured from neuronal culture supernatants using a Promega Glutamate-Glo assay kit (Promega, # J7021, Table S1), according to the manufacturer’s instructions. Measurements were normalized to total protein content in the lysates.

### Cell homogenate

Cells were lysed in isotonic buffer (0.32 M sucrose, 20 mM HEPES (free acid), 150 mM NaCl, pH adjusted to 7.4 with NaOH) supplemented with EGTA-free protease inhibitor cocktail (Calbiochem, # 539134-1ML; 1:1000) and phosphatase inhibitors PhosSTOP^TM^ (Sigma-Roche, #4906845001). The buffer did not use any surfactants so as to maintain the native state of proteins in their self-assembly and/or aggregation phenotypes.

### ELISA

Detailed protocols for sandwich ELISA and signal amplification are as provided previously (Kapadia *et al*, 2025; Kapadia & Hafner, 2025). Briefly, equal amounts of protein from cellular homogenates were added to antibody precoated Nunc® MaxiSorp™ 384 well plates (Sigma Aldrich, #P6366) and incubated overnight at 4°C. Plates were washed, blocked with 1% BSA in PBS (2 h, RT), and incubated with primary detection antibodies (Table S2, 2 h at 37°C or overnight at 4°C) followed by HRP-conjugated secondary antibodies (3 h at 37°C). Detection was performed using TMB substrate (ThermoFischer, #34029), and absorbance was measured at 450 nm with background correction at 620 nm. Samples were analyzed in duplicates or triplicates.

### Western blotting

Equal amounts of protein (10–20 μg), determined using the Pierce BCA assay (ThermoFisher Scientific, #23225); were loaded per lane unless stated otherwise. For SDS-gel electrophoresis, samples were boiled at 95°C for 5 min. Samples for native gel electrophoresis were not boiled. Briefly, proteins were separated on 4–20% Bis-Tris gels (Gibco #) using an NuPAGE™ MES SDS Running Buffer (Invitrogen, #NP0002) or NativePAGE™ Running Buffer (Invitrogen, #BN2001) and transferred to 0.2 μm nitrocellulose (Amersham, #1060001) or PVDF (Amersham, #10600021) membranes. Membrane was blocked with 5% skimmed milk (Cal Roth, #T145-1) in TBS-T containing 0.1% Tween 20 (ThermoFischer Scientific, #J20605.AP)) for 1 h, RT; and incubated with primary antibodies (overnight at 4°C, Table S2), followed by secondary antibodies for 2 h at RT (Table S2). Detection was performed using ECL (Bio-Rad, Germany) or Li-COR fluorescent imaging systems (LiCOR Biosciences, Germany). Signals were normalized to control conditions within each experiment.

### Single-molecule pull-down (SIMPull)

SIMPull assays were performed as described earlier (Kapadia and Hafner, 2025), using antibody-functionalized glass surfaces. APP-derived species were captured using 4G8 antibody and detected using A11 or vesicle markers. Images were acquired using TIRF microscopy and analyzed for puncta density and colocalisation using ImageJ FiJi analysis module.

### GPMV preparation, labelling and imaging

Giant plasma membrane vesicles (GPMVs) were prepared from CHO cells using chemical induction as previously described (Sezgin et al., 2012; Yosibash et al., 2026). Cells were cultured in 10 cm dishes (Corning, Sigma Aldrich, # CLS430167) attaining about 80-90% confluency. To induce vesiculation, the cells were incubated in vesiculation buffer (10 mM HEPES (pH 7.4), 150 mM, NaCl_2_ and 2 mM CaCl₂ supplemented with 2 mM N-ethylmaleimide (NEM), 25 mM paraformaldehyde (PFA), 2 mM dithiothreitol (DTT), and 1mM ATP) overnight at 37_°_C. Next day, the GPMVs were then transferred to the imaging chanmbers and incubated with the dye mixture for 15-20 min. GPMVs were live-labelled with lipid probes including CellTracker Green BODIPY (ThermoFischer, #C2102), Fillipin (Sigma-Aldrich, #SAE0087), Wheat Germ Agglutinin (WGA)- Alexa Fluor Plus 568 conjugate (ThermoFischer, #W56133), and DiIC12(3) perchlorate (AAT Bioquest, #22035). APP/C99 was live-labelled with 4G8-Alexa 488 conjugated antibody. Background was slightly higher as dyes remained in solution during imaging. Thus, care was taken to dilute the dye solution (1:1, best possible alternative) after the GPMVs settled on the coverslips before image acquisition. Imaging was performed using Nikon TIRF microscopy on widefield settings.

### Amplex Red Cholesterol assay

Cellular and GPMV-associated cholesterol levels were quantified using the Amplex™ Red Cholesterol Assay Kit (Thermo Fisher Scientific, #A12216; Table S1), according to the manufacturer’s instructions and previous optimisation (Kapadia et al., 2025). Equal amounts of protein from cell lysates or isolated GPMV fractions were loaded into 96-well plates and incubated with freshly prepared Amplex Red working solution (containing cholesterol oxidase, cholesterol esterase, Amplex Red and horseradish peroxidase) for 30 min at 37°C protected from light. Fluorescence intensity of the produced resorufin was measured using a plate reader (Ex: ∼560 nm, Em: ∼590 nm), and cholesterol levels were quantified relative to a cholesterol standard curve (kit standard) and normalized to total protein content per sample for independent experiments, respectively.

### Synaptic vesicle isolation

Synaptic vesicles were isolated from rat brain homogenates using differential centrifugation as described earlier (Kapadia and Hafner, 2025). In short, synaptosomes and synaptic vesicles were enriched from the forebrains of 1-year-old wild-type rats. To favour the isolation of synaptosomes with presynaptic compartments, we used the previously described protocol, (Hafner et al., 2019; Kapadia et al., 2025; Kapadia and Hafner, 2025; Luquet et al., 2017). Forebrain from a rat hemibrain homogenized in 2 ml of ice-cold homogenization buffer [0.32 M sucrose (Merck, #S9378-1KG), 4 mM HEPES (Sigma Aldrich, #H4034-100G; pH adjusted to 7.4), EGTA-free protease inhibitor cocktail (Calbiochem, # 539134-1ML; 1:1000), by using a 2-ml glass-Teflon homogenizer with 15-17 gentle strokes. An additional 2 ml of buffer was added and centrifuged at 1000 × g for 8 min at 4°C. The supernatant (S1) was centrifuged against 12,500 × g for 13 min at 4°C. The synaptoneurosome-enriched pellet (P2) was then resuspended in 1 ml of homogenization buffer. This fraction was finally layered on top of a two-step sucrose density gradient (5 ml of 1.2 M sucrose and 5 ml of 0.8 M sucrose, 4 mM HEPES, and EGTA-free protease inhibitor cocktail, as described above). The gradient was ultracentrifuged at 40,000 × g for 70 min at 4°C. Synaptosome fractions were recovered through the tube wall, at the interface of 0.8 and 1.2 M sucrose, by using a syringe to minimize contamination with lighter fractions enriched in myelin. The resulting fraction is referred to as synaptosomes (SYN). Next, SYN fraction was homogenised in SynPer buffer and reloaded on the two-step sucrose density gradient and ultracentrifuged for 120,000 × g for 2 h at 4°C. Again, synaptic vesicle fractions were recovered through the tube wall, at the interface of 0.8 and 1.2 M sucrose, by using a syringe to minimize contamination. Aliquots were prepared and stored at -80°C, until further use. It is recommended to prepare smaller aliquots as repeated freeze thawing of the SV fractions reduces its viability for experiments. Isolated SV aliquots were labelled with SynaptoRed C2 (Sigma Aldrich, #S6689-5MG, 16 nM in SynPer buffer) prior to use in tethering assays. Again here, it is recommended to use freshly prepared material on each day of the experiment.

### Synaptic vesicle recruitment assays using SIMPull

Extended SV SIMPull assays were performed as described earlier (Kapadia & Hafner, 2025), using antibody-functionalized glass surfaces. APP-derived species were captured using 4G8 antibody and detected using APP-C-terminal antibody clone: C1.6.1 wherever required. For vesicle tethering assays, purified synaptic vesicles labelled with SynaptoRed were incubated with captured APP species. Images were acquired using TIRF microscopy and analyzed for puncta density.

### Membrane sheet preparation

Membrane sheets were generated as previously described (Perez et al., 2006), from CHO cells by brief sonication, exposing the cytosolic face of the plasma membrane. Briefly, cells plated on 25 mm glass coverslips and were used with 70-80% confluency. The cytosolic buffer consisted of 25 mM HEPES, 1.5 mM MgCl₂, 10 mM KCl, and 1 mM EDTA in distilled water. The solution was adjusted to pH 7.4 prior to volume finalization. Immediately before experimental use, the buffer was supplemented with 1 mM dithiothreitol (DTT) and a 1× protease inhibitor cocktail to minimize protein degradation. All buffer solutions were kept ice-cold throughout the isolation procedure. Coverslips were incubated in cytosolic buffer in a 50 ml glass beaker and subjected to probe sonication (35 mW power, 3 cycles) while maintaining the entire setup on ice. Coverslips were then assembles into the imaging chamber and incubated with nuclear (DAPI) or membrane dyes, such as CellTracker Green BODIPY (ThermoFischer, #C2102), Phalloidin-647 conjugate (Sigma-Aldrich, #SAE0087), Wheat Germ Agglutinin (WGA)- Alexa Fluor Plus 568 conjugate (ThermoFischer, #W56133) to ensure optimal preparation of membrane sheets and the removal of cytosolic /nuclear compartments.

### Synaptic vesicle recruitment assays using functionalised membrane sheets

Sheets were incubated with synaptic vesicles fluorescently labelled with SynaptoRed for 20 min. Using an in-house flow through set-up attached to a syringe pump, washes were performed as described earlier (Kapadia & Hafner, 2025). Vesicle tethering was quantified relative to membrane area by creating 2 masks, mask1 = phalloidin positive area, and mask2 = 4G8 positive area. Tethered SR-SV counted using FiJi ImageJ > analyse particles tool. Particle size cutoff was limited to 0.5 micrometer, particles bigger than that were eliminated. It is important to keep in mind that this could slightly underrepresent the data quantified for C99^YENPTY^ and C99^ANAETA^ variants. Quantification depicts % change in bound SR-SV puncta were computed based on the formula [(mask 2/ mask 1) x 100].

### Microscopy and image processing

Confocal images were acquired using Nikon SoRa spinning disk super-resolution microscope (Imaging lasers:405, 445, 488, 561, 638 nm equipped with Emission filters: UV, green, yellow, red, far-red) using a 40-x oil immersion Nikon objective (SoRa magnifiers 2.8x) with N/A 1.3 (working distance: 200 μm), or Nikon TIRF microscope (TIRF-mode with lasers: excitations: 488 nm, 561 nm, 640 nm) or 100-x oil immersion Nikon objective with N/A 1.49 (working distance: 160 μm). Imaging parameters, laser intensity and set-up were kept consistent across conditions. Images were analysed and processed minimally (if any) using FiJi ImageJ software.

### Statistical analysis

Data were analyzed using GraphPad Prism (v10.4.1). Results are presented as mean ± SEM. ‘N’ represents independent biological replicates, and ‘n’ represents technical replicates. Statistical tests are indicated respectively in figure legends. All parametric and non-parametric tests have been performed using the in-built default settings of the module in the software, without modifications, until mentioned otherwise; and significance thresholds were defined as: ns (*p* > 0.05), \**p* ≤ 0.05, \*\**p* ≤ 0.01, \*\*\**p* ≤ 0.001.

## Acknowledgements

We would like to thank all our lab members and Dr. Marijn Kuijpers (MK, Radboud University, Nijmegen, Netherlands) for fruitful discussions and helpful feedback. APP/C99 plasmids were a gift from Dennis Selkoe & Tracy Young-Pearse. Synapto-pHluorin was a gift from Franck Polleux (Addgene plasmid #168510) to MK. Prof. Jochen Walter (University of Bonn, Germany) shared APP and Aβ antibodies. Dr. Eric Jansen and Prof. Corette Wierenga (Radboud University, Nijmegen) shared CHO cell lines. Dr. Jelle Postma 9general instrumentation, Radboud University, Nijmegen) assisted with the TIRF microscopy set-up. This study was supported by the Donders Institute for Brain, Cognition and Behavior and Faculty of Science, Radboud University Nijmegen Netherlands. Schematics were created using Biorender.com.

## Author contributions

Conceptualization: AK

Methodology: AK

Investigation: AK, EC, SV

Visualization: AK, EC

Funding acquisition: AH

Project administration: AH

Supervision: AK, AH

Writing – original draft: AK

Writing – review & editing: AK, EC, SV, AH

## Data and materials availability

All experimental data in the main text or the supplementary materials will be made available at the Donders data repository. All numerical values pertaining to the graphical illustrations in the figures have been summarised in the Supplementary Data file 1.

## Supplementary Materials

Figures S1 to S6

Tables S1

Data file 1

