## Supplementary Material file for "APP-CTFβ/C99 oligomers drive synaptic vesicle tethering through C-terminal interactions"

**Supplementary data associated with the manuscript contains:**

Tables S1 and S2

Figures S1 to S5

Data file 1

**Table S1: List of materials and reagents**

| Reagent / Material | Supplier | Catalog Number |
|---|---|---|
| ***DNA Constructs*** | | |
| APP695 (APP-FL) [Human APP] | Addgene | #30137 |
| APP-CTFβ (C99) / C99^YENPTY [YENPTY (non-mutated)] | Addgene | #30146 |
| C99^ANAETA [YENPTY --> ANAETA (mutant)] | Addgene | #30144 |
| C99^ΔCT [ C-terminal deletion] | Addgene | #30143 |
| Synaptophysin-pHluorin Reporter construct | Addgene | #168510 |
| ***Cell Line/Biological material*** | | |
| CHO cells | ATCC | CCL-61 |
| Primary cortical neurons | Janvier (E18 rat embryos) | |
| Synaptic vesicles | Janvier Adult Rat | |
| ***Cell Culture Medium, Supplements and Reagents*** | | |
| Basal Medium Eagle | Thermo Fisher | #21010046 |
| BrainPhys™ medium | STEMCELL | #05790 |
| NeuroCult™ SM1 | STEMCELL | #05711 |
| B27 Plus | Gibco | #A3582801 |
| Penicillin-Streptomycin | Gibco | #15140122 |
| NEAA | Thermo Fisher | #11140050 |
| Sodium pyruvate | Thermo Fisher | #11360070 |
| Ham's F-12 | Gibco | #11765047 |
| Fetal bovine serum | Thermo Fisher | #A5209501 |
| Poly-D-lysine | Sigma | #A-003-E |
| Glass coverslips | VWR | #631-1577/79/84 |
| Lipofectamine 2000 | Thermo Fisher | #11668027 |
| G418 | InvivoGen | #ant-gn-1 |
| ***Pharmacological inhibitors*** | | |
| DAPT (GSI) | Cayman | #13197-5 |
| Tetrodotoxin (TTX) | HelloBio | #HB1035 |
| ***Miscellaneous material and reagents*** | | |
| Paraformaldehyde | Thermo Fisher | #A11313.36 |
| Sucrose | Sigma | #S9378-1KG |
| PBS | VWR | #A0965.9010 |
| Triton X-100 | Merck | #T8787-250ML |
| BSA | Carl Roth | #8076-4 |
| Normal horse serum | Gibco | #26050088 |
| Moviol 4-88 | Carl Roth | #0713.1 |
| MaxiSorp plates | Sigma | #P6366 |

| | | |
|---|---|---|
| TMB | Thermo Fisher | #34029 |
| BCA protein assay | Thermo Fisher | #23225 |
| Bis-Tris gels | Gibco/Bio-Rad | – |
| MES running buffer | Invitrogen | #NP0002 |
| NativePAGE buffer | Invitrogen | #BN2001 |
| Nitrocellulose 0.2µm | Amersham | #1060001 |
| PVDF 0.2µm | Amersham | #10600021 |
| Skim milk powder | Carl Roth | #T145-1 |
| Tween-20 | Thermo Fisher | #J20605.AP |
| Protease cocktail | Calbiochem | #539134 |
| PhosSTOP™ | Sigma Roche | #4906845001 |
| Promega Glutamate-Glo assay kit | Promega | #J7021 |
| Amplex™ Red Cholesterol Assay Kit | Thermo Fisher Scientific | #A12216 |
| ***Dyes for staining*** | | |
| SynaptoRed C2 (Synaptic vesicles) | Sigma | #S6689-5MG |
| BODIPY (Lipid probe) | Thermo Fisher | #C2102 |
| Filipin (Cholesterol) | Sigma | #SAE0087 |
| WGA-Alexa 568 (Membrane marker) | Thermo Fisher | #W56133 |
| DiIC12(3) (raft marker) | AAT Bioquest | #22035 |
| ***Instrumentation and Software*** | | |
| SoRa Spinning disk microscope | Nikon Technologies, Japan | |
| TIRF microscope | Nikon Technologies, Japan | |
| Imaging software | Nikon Technologies, Japan | |
| Odessey M | LiCOR, Germany | |
| ECL imager | BioRAD, Germany | |
| Image Studio Lite (WB processing and Image generation) | LiCOR, Germany | |
| GraphPad Prism (Statistics) | GraphPad | v10.4.1 |
| Fiji/ImageJ (Image analysis) | NIH | – |

**Table S2: List of antibodies used in this study.**

| Target of interest and Specificity (Antibody clone) | Make (Cat. No.) | WB | ICC | ELISA / SIM-Pull |
|---|---|---|---|---|
| ***Amyloid precursor protein (APP), APP C-terminal fragments and Amyloid β (Aβ)*** | | | | |
| N-terminal specific | Sigma Aldrich (SAB4200536) | | 1:250 | 1:100 |
| N-terminal fragment specific (clone: 3207) | In house generated (Walter lab, Bonn) (Eurogentec) | | 1:500 | 1:200 |
| APP-Aβ mid region- specific (clone: 4G8) | Biolegend (800708) | 1:1000 | 1:500 | 1:200 |
| 4G8 -Biotin conjugated | Biolegend (800704) | | | 1:1000 |
| 4G8 -Alexa 488 conjugated | Biolegend (800714) | | 1:1000 | |
| Aβ1-x (clone: 82E1) | IBL Int. (JP10323) | 1:500 | 1:200 | |
| 82E1-Biotin conjugated | IBL Int. (JP10326) | | | 1:1000 |
| APP C-terminal fragment specific (clone: C1/6.1) | Biolegend (802801) | 1:1000 | 1:1000 | 1:1000 |
| Aβ (clone: 2964) | In house generated (Walter lab, Bonn) (Eurogentec) | | 1:1000 | |
| Oligomer A11 Polyclonal Antibody | ThermoFischer (AHB0052) | | | |
| ***Cellular markers*** | | | | |
| *Excitatory synapse:* Vesicular Glutamate Transporter (vGLUT1) | SYSY (135 304) | 1:1000 | 1:2000 | 1:1000 |
| *Pan-synaptic marker:* Synapsin | SYSY (106 011 and 106 004) | | | 1:500 |
| *Synaptotagmin-1 (Syt1):* C-terminal cytoplasmic domain | SYSY (105 008 and 105 011), DSHB (mAB30-asv30) | 1:1000 | | 1:500 1:1000 (1:1) |
| *Synaptobrevins (Syb)* | SYSY (104 002) | 1:1000 | | 1:500 1:1000 (1:1) |
| *Vesicle-associated membrane protein- 1/2/3 (VAMPs-1/2/3)* | SYSY (104 102) | 1:1000 | 1:500 | 1:1000 (1:1) |
| *Somatodendritic marker:* Microtubule protein (MAP2) | SYSY (188004 and 188 006) | | 1:1000 | |
| *Axon marker:* Neurofilament-L | SYSY (171 002) | | 1:500 | |
| *Beta-tubulin (β3TUBB)* | SYSY (302 303) | 1:1000 | | |
| *Nucleus* stain: 4′,6-diamidino-2-phenylindole (DAPI) | TFS (D1306) | | | |

| | | | | |
|---|---|---|---|---|
| ***Fluorescent conjugated secondary antibodies*** | | | | |
| Goat anti-chicken IgG 405 | Abcam (ab175674) | | 1:500 /1000 | |
| Goat Anti-Chicken IgY H&L A488 | Abcam (ab150169) | | | |
| Goat anti-Mouse IgG A405 | Ozyme (BTM20080-1MG) | | | |
| Goat anti-Mouse IgG A488 | Abcam (ab150113) | | | |
| Goat anti-Mouse IgM (Heavy chain) A568 | Invitrogen (A21043) | | | |
| Goat anti-Mouse IgG A647 | Abcam (ab150115) | | | |
| Goat anti-Mouse IgG Atto647 | Sigma Aldrich (50185-1ML-F) | | | |
| Goat anti-Rabbit IgG A488 | Abcam (ab150077) | | | |
| Goat anti-Rabbit IgG A568 | Abcam (ab175471) | | | |
| Goat anti-Rabbit IgG A568 | Ozyme (BTM20102-1MG) | | | |
| Goat anti-Guinea Pig IgG A488 | Abcam (ab150185) | | | |
| Goat anti-Guinea Pig IgG A647 | Abcam (ab150187) | | | |
| Streptavidin Alexa 568 Conjugate | TFS (S11226) | | 1:1000 | |
| Streptavidin, Alexa Fluor™ 647 conjugate | Biolegend (405237) | | | |
| ***HRP conjugated secondary antibodies*** | | | | |
| Goat anti-Mouse IgG (H+L) Secondary Antibody, HRP | TFS (32430) | 1:2500/5000 | 1:1000 | |
| Goat anti-Rabbit IgG (H+L) Secondary Antibody, HRP | TFS (31460) | | | |
| Goat anti-Guinea Pig IgG (H+L) Secondary Antibody, HRP | TFS (A18769) | | | |
| Streptavidin HRP | Biolegend (1474) TFS (N100) | 1:2500 | | 1:1000/2500 |

*WB, western blotting; ICC, immunocytochemistry; ELISA, Enzyme linked immunosorbent assay; SIM-Pull, single molecule pull-down imaging assay. For APP, C99 and Aβ-specific antibody epitope-specificity please see SI, Scheme 1.*

**Scheme 1: Overview of APP-FL, APP-CTFβ, and Aβ species highlighting epitope specificity and detection range of the antibodies used in this study.**

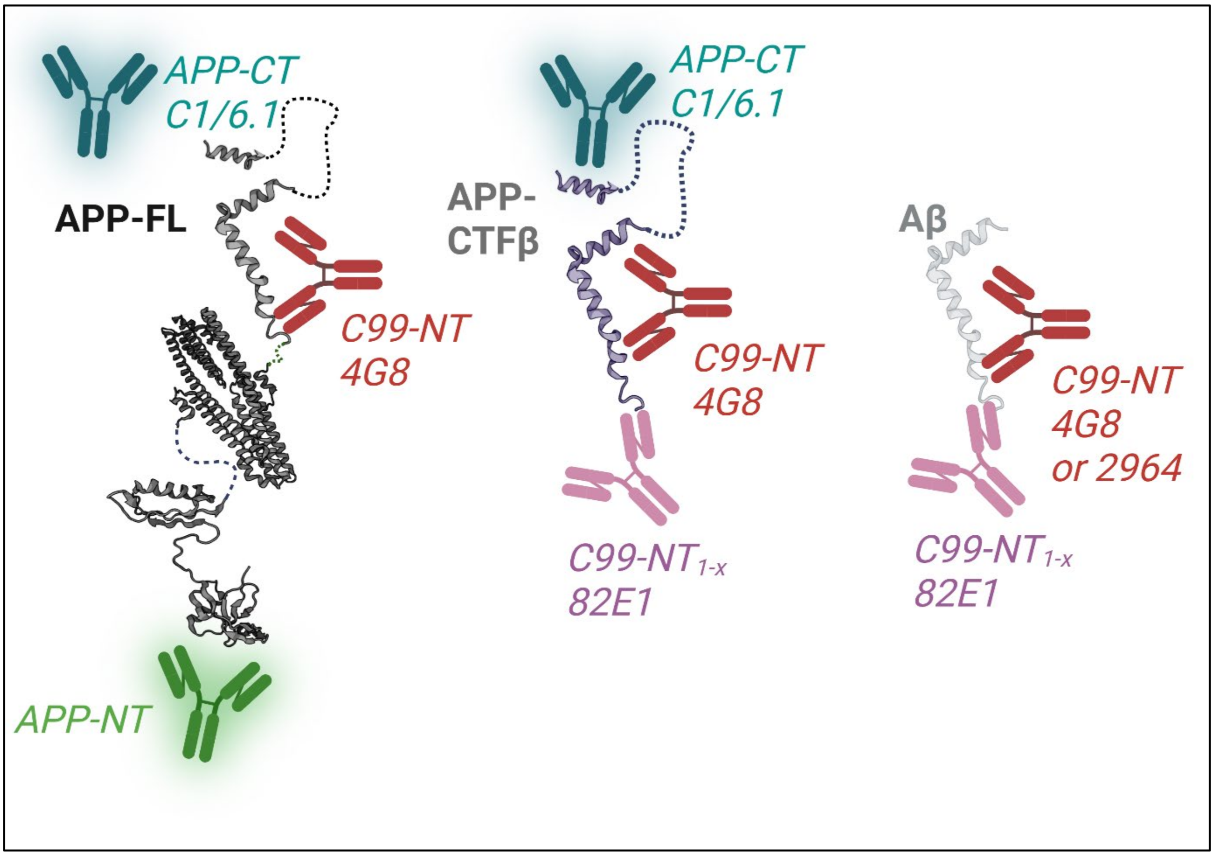

### SI Figures and legends:

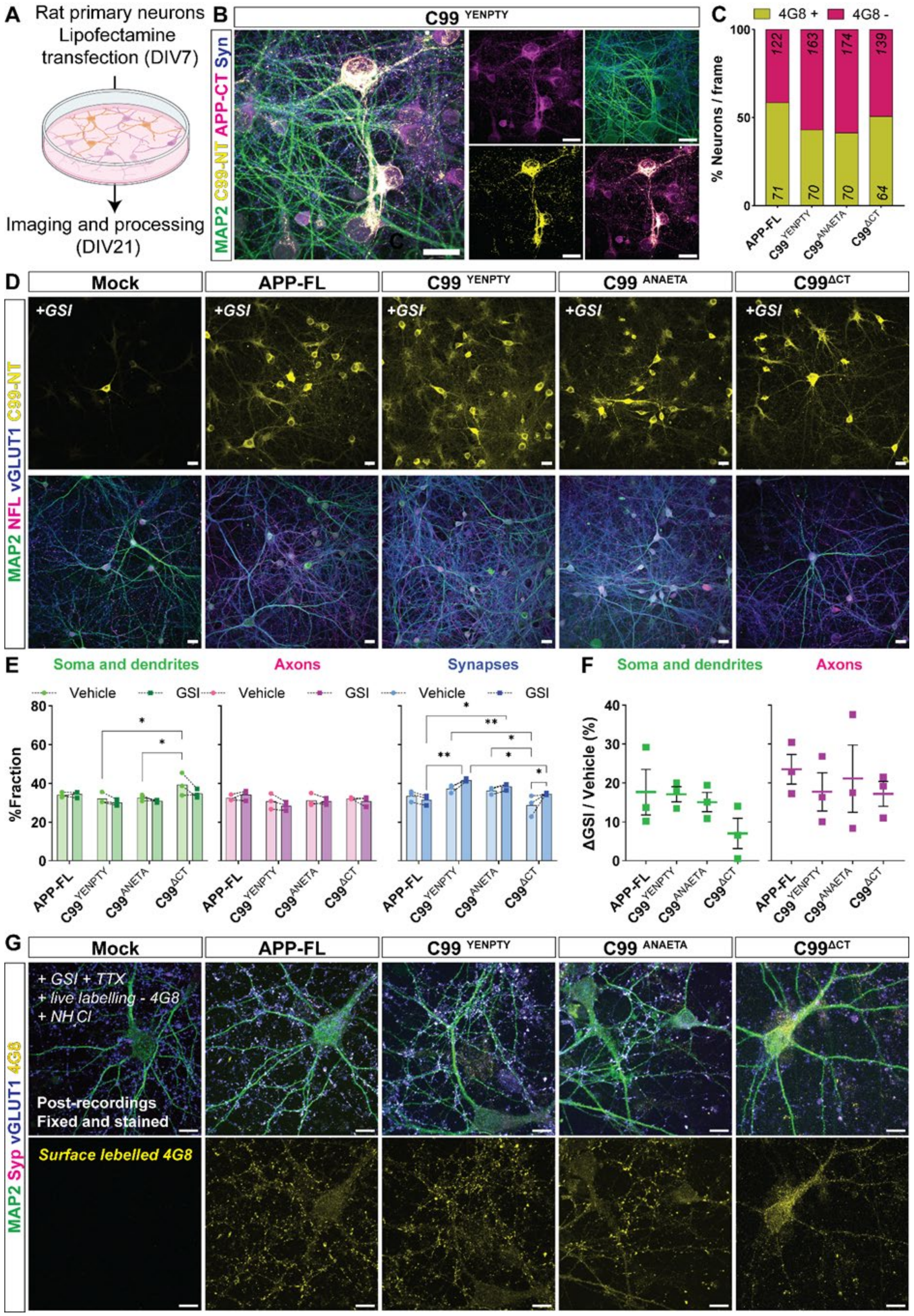


**Figure S1. APP-CTFβ expression, distribution, and functional validation in primary neurons.**

**(A)** Schematic of experimental workflow for transient transfection of rat primary cortical neurons (DIV7) followed by pharmacological treatment, imaging and subsequent analysis at DIV21.

**(B)** Representative immunocytochemistry images of neurons expressing human APP-CTFβ (C99$^{YENPTY}$), co-stained with 4G8 (*yellow*), MAP2 (*green*), global APP C-terminal species (antibody clone; C1/6.1, *magenta*), and Syn1/2 (*blue*). Panels to the right show colocalization between individual channels. Scale bars, 10 µm**.**

**(C)** Quantification of transfected (4G8-positive) and non-transfected (4G8-negative) neurons per imaging field across APP/C99 variants, indicating proportion of 4G8-positive and 4G8-negative neurons per imaging field; across the different APP/C99 variants.

**(D)** Representative images of neurons expressing APP-FL and C99 variants following γ-secretase inhibition (GSI; DAPT- 10 µM, 4 h) treatment. Neurons were stained for 4G8 (*yellow*) alongside MAP2 (*green*), NFL (*magenta*), and Syn1/2 (*blue*). Scale bars, 10 µm.

**(E)** Quantification showing the percentage distribution of APP/C99 signals across somato-dendritic (*green*), axonal (*magenta*), and synaptic compartments (*blue*) under vehicle (*lighter bars*) and GSI treated (*darker bars*) conditions, illustrating preferential accumulation of APP/C99 species within synaptic compartments following γ-secretase inhibition. Individual points represent average from each independent experiment; lines depict respective changes post-treatment in each individual experiment; bars, mean; error bars, S.D.; N = 3 independent transfected cultures.

**(F)** Scatter plot depicting percentage change (Δ) of APP/C99 overlap with MAP2-positive somato-dendritic compartments (*green*) and NFL-positive axons (*magenta*) following GSI treatment relative to vehicle (untreated) controls. ΔGSI/Vehicle (%) = $[(MOC_{GSI} - MOC_{Vehicle}) / MOC_{Vehicle}] \times 100$. Values are expressed as percentage change relative to the corresponding vehicle-treated condition. Individual points represent average from each independent experiment; bold line, mean; error bars, S.D.; N = 3 independent transfected cultures.

(G) Representative images of neurons following synaptophysin-pHluorin recordings, fixed and stained for MAP2 (*green*), synaptophysin (Syp, *magenta*), and vGLUT1 (*blue*). Bottom panels show surface-labelled APP/C99 species detected using 4G8, confirming that analyzed pHluorin-positive neurites correspond to APP/C99-expressing processes. Scale bars, 10 µm.

Statistical significance was determined using one-way ANOVA with post hoc multiple-comparison testing as indicated. $^{ns}$ $p > 0.05$, * $p < 0.05$, ** $p < 0.01$, *** $p < 0.001$.

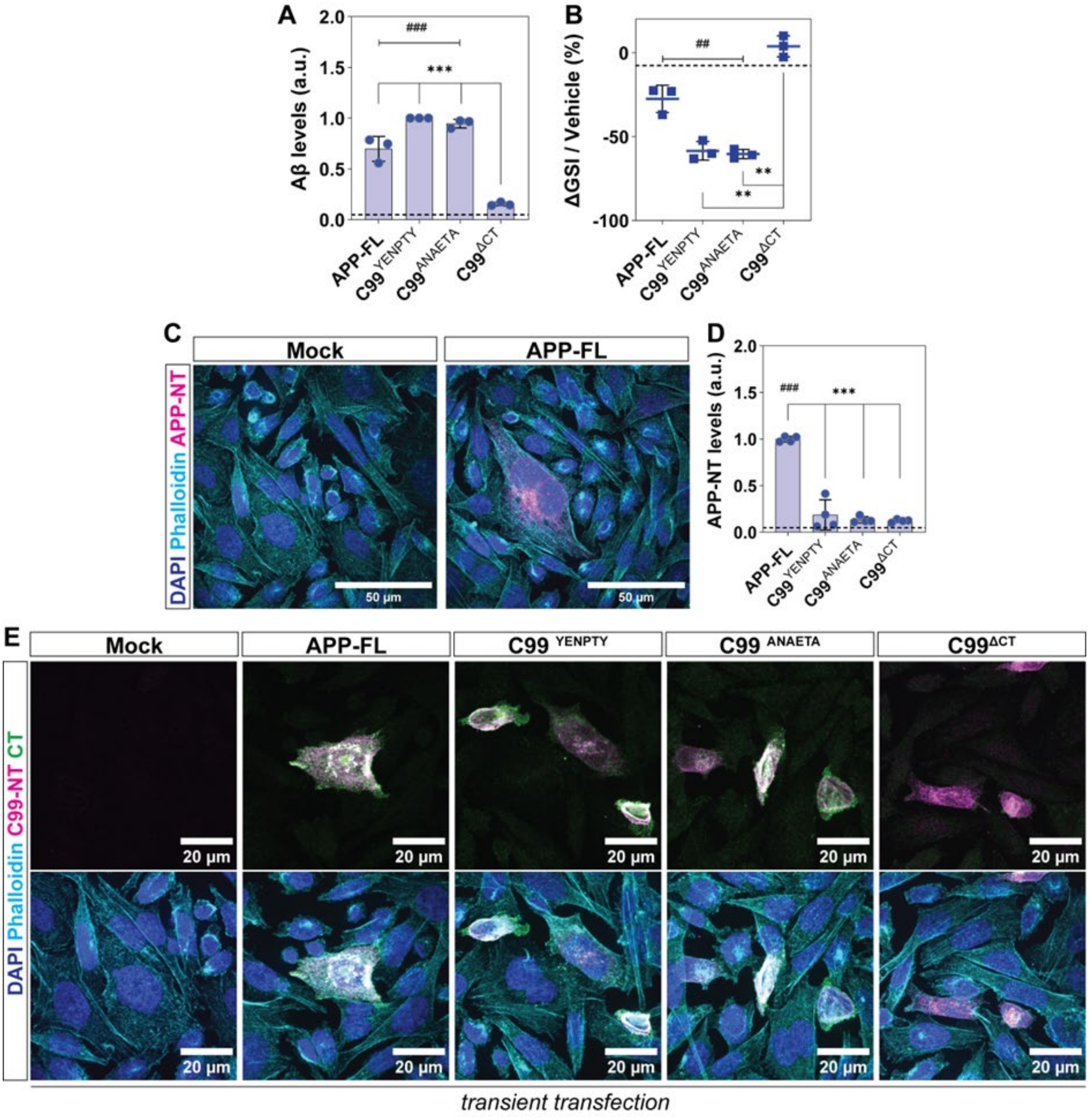


**Figure S2. γ-secretase-dependent changes in Aβ production in CHO cells and validation of APP/C99 expression in transiently transfected cells.**

**(A)** ELISA analysis depicting extracellularly secreted Aβ species (antibody clone: 2964) in conditioned media from APP-FL- and C99-expressing stable CHO cells under basal conditions. Individual points represent average from each experiment normalised to C99$^{YENPTY}$ condition within each biological replicate; bars, mean; error bars, S.D., dotted line, mock; N = 3 independent cultures.

**(B)** Scatter blots depicting percentage change (Δ) of extracellular Aβ levels (antibody clone: 2964; ELISA), following GSI treatment relative to the corresponding vehicle-treated condition. ΔGSI/Vehicle (%) = [(Signal$_{GSI}$ - Signal$_{Vehicle}$) / Signal$_{Vehicle}$] ×100. Values are expressed as percentage change relative to the corresponding vehicle-treated condition. Individual points represent average from each independent experiment; bold line, mean; error bars, S.D., dotted line, mock; N = 3 independent transfected cultures.

**(C)** Representative immunocytochemistry images validating selective detection of APP-FL using an APP N-terminal ectodomain antibody (APP-NT, *magenta*). Cells were counterstained with phalloidin (*cyan*) and DAPI (*blue*). Scale bars, 50 µm.

**(D)** Bar plots depict quantification of intracellular APP-FL (antibody clone; APP-NT, ELISA), from APP-FL- and C99-expressing stable CHO cells under basal conditions. As expected, the APP N-terminal ectodomain was detected exclusively in APP-FL-expressing cells and was absent from all C99 variants. Individual points represent average from each experiment normalised to APP-FL condition within each biological replicate; bars, mean; error bars, S.D., dotted line, mock; N = 3 independent cultures.

**(E)** Representative immunocytochemistry images of transiently transfected CHO cells expressing APP-FL or C99 variants. Cells were stained with 4G8 (*magenta*), APP C-terminal antibody C1/6.1 (*green*), phalloidin (*cyan*), and DAPI (*blue*). Scale bars, 20 µm. Transiently transfected cells were subsequently used for membrane sheet generation and vesicle tethering experiments (Fig. 4, Fig. 5 and Fig. S6). Data is representative from 2 independent experiments.

Statistical significance was determined using one-way ANOVA with post hoc multiple-comparison testing. Comparisons among APP/C99 variants are denoted by asterisks (*), whereas comparisons relative to mock controls are denoted by hash symbols ($^{\#}$). $^{ns}$ $p > 0.05$, $^{*/\#}$ $p < 0.05$, $^{**/\#\#}$ $p < 0.01$, $^{***/\#\#\#}$ $p < 0.001$.

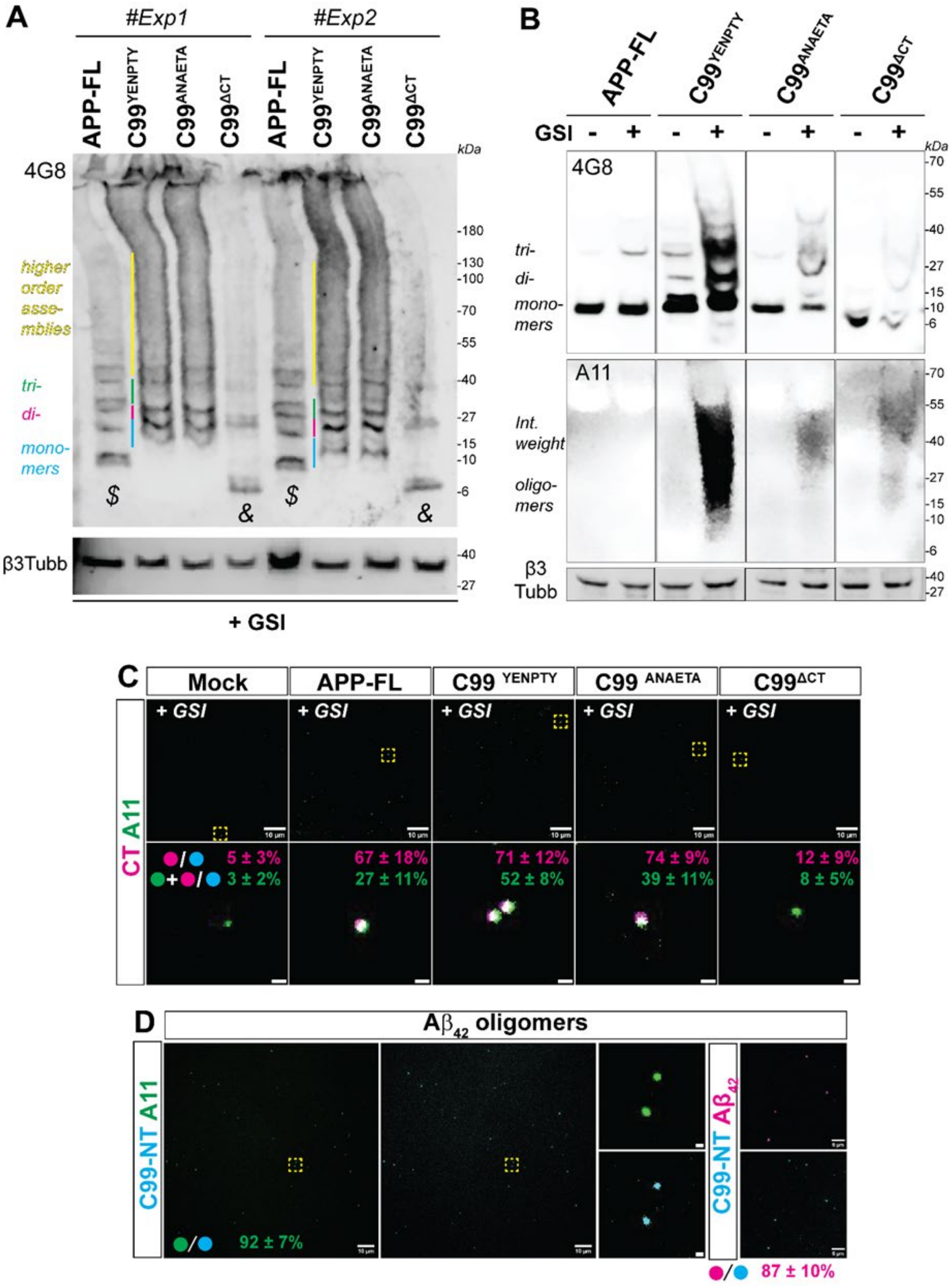


**Figure S3. Orthogonal validation of APP-CTFβ oligomer formation following γ-secretase inhibition**

**(A)** Western immunoblots showing accumulation of C99 species and higher molecular weight 4G8-immunoreactive assemblies in stable CHO cells expressing APP-FL or C99 variants following γ-secretase inhibition (+GSI). β3-tubulin (β3-tubb) serves as loading control. Approximate positions of monomeric (*cyan*), dimeric (*magenta*), trimeric (*green*), and higher-order assemblies (*yellow*) are indicated. Two independent experiments demonstrate reproducible accumulation of higher molecular weight APP-derived species, particularly in $C99^{YENPTY}$ and $C99^{ANAETA}$ expressing cells (Fig. 2h and 3A). $, APP-derived CTFs signals; & $C99^{\Delta CT}$ species.

**(B)** Native western blot analysis of stable CHO cell lines expressing APP-FL or C99 variants under vehicle (-GSI) and γ-secretase inhibited (+GSI) conditions. 4G8 immunoreactivity reveals accumulation of monomeric and higher molecular weight APP-derived species following γ-secretase inhibition. A11 immunoreactivity confirms the presence of oligomeric assemblies enriched following γ-secretase inhibition, most prominently in $C99^{YENPTY}$ and $C99^{ANAETA}$ expressing cells. β3-tubulin served as loading control.

**(C)** Representative SIMPull images showing capture of APP-derived species using a 4G8 antibody and simultaneous detection using C1/6.1 antibody (CT) and the oligomer-specific A11 antibody under +GSI conditions. Insets show magnified puncta. A11-positive puncta lacking corresponding C1/6.1 immunoreactivity were observed in mock and $C99^{\Delta CT}$ controls and were excluded from colocalization analyses. Data represent mean ± S.D. from N = 3 independent experiments. Scale bars, 10 µm (*top*), 0.2 µm.

**(D)** Positive-control SIMPull experiments using pre-assembled Aβ42 oligomers using a similar SIM-Pull strategy as described in panel C. Aβ42 oligomers served as a positive control for A11 immunoreactivity and were used for normalization of oligomer-specific ELISA and SIMPull analyses. Since Aβ lacks the C-terminal epitope recognised by C1/6.1, the Aβ42-specific antibody BAP-15 was used. N = 2 independent experiments. Scale bars, 10 µm, zoomed panels, 0.2 µm.

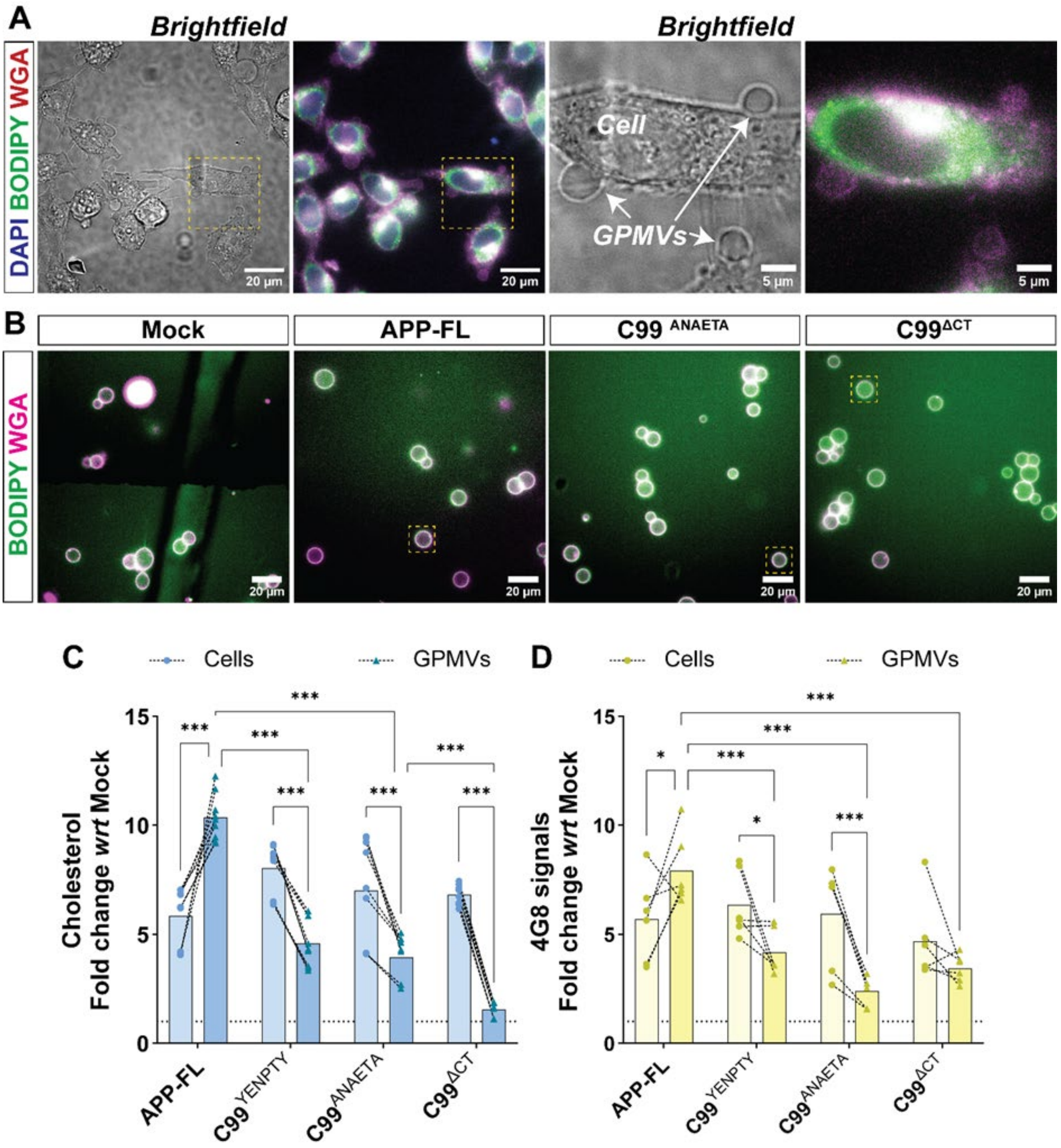


**Figure S4. APP-CTFβ-associated changes in GPMV membranes.**

**(A)** Representative brightfield and fluorescence images illustrating the formation of giant plasma membrane vesicles (GPMVs) from CHO cells. GPMVs were labelled with BODIPY (lipids, green), wheat germ agglutinin (WGA; membrane glycoproteins, magenta), and DAPI (blue). Zoomed in panels highlight morphological differences between intact cells and detached GPMVs. Data are representative of N = 2 independent experiments. Scale bars, 20 µm; zoomed panels, 5 µm.

**(B)** Representative images of GPMVs derived from mock, APP-FL, $C99^{ANAETA}$, and $C99^{\Delta CT}$ expressing cells, co-labelled with BODIPY (*green*) and WGA (*magenta*). Scale bars, 20 µm. N = 4 independent experiments.

**(C)** Quantification of cholesterol levels in whole-cell lysates (circles) and corresponding GPMV fractions (triangles) measured using the Amplex Red cholesterol assay. Values are expressed as fold change relative to mock controls. Individual symbols represent independent experiments connected by dashed lines; n = 6 technical replicates, N = 3 independent experiments.

**(D)** Quantification of APP/C99 species detected with 4G8 antibody in whole-cell lysates (circles) and corresponding GPMV fractions (triangles). Values are expressed as fold change relative to mock controls. Individual symbols represent independent experiments connected by dashed lines; n = 6 technical replicates, N = 3 independent experiments.

Statistical significance was determined using two-way ANOVA with post hoc multiple-comparison testing. * $p < 0.05$, ** $p < 0.01$, *** $p < 0.001$.

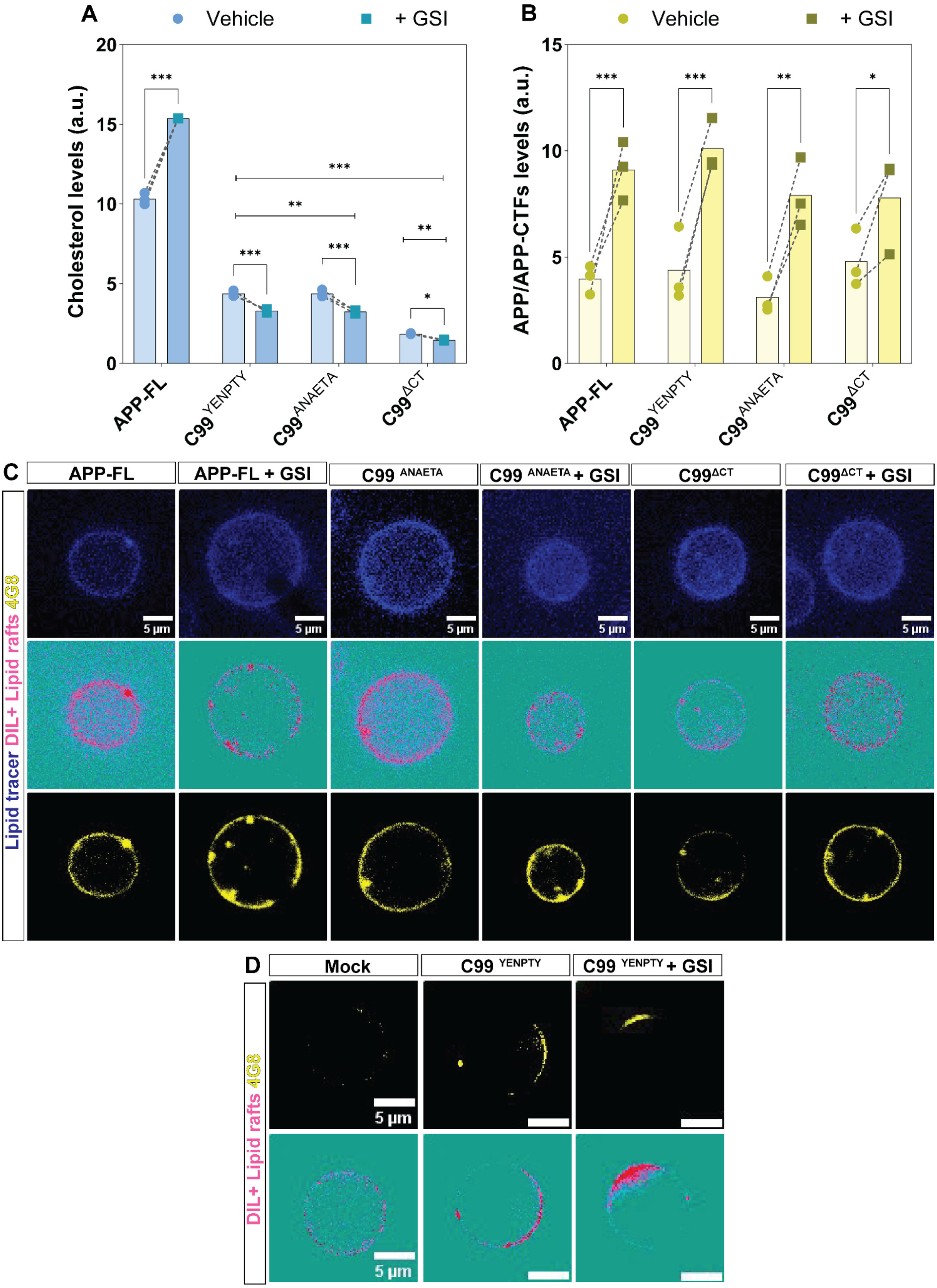


**Figure S5: γ-Secretase inhibition alters APP-CTFβ partitioning and membrane lipid organization in GPMVs**

**(A)** Quantification of APP/APP-CTF species in GPMVs derived from APP-FL- and C99-expressing CHO cells under vehicle (circles) and γ-secretase inhibited (+GSI; squares) conditions, measured by ELISA. Values are expressed as fold change relative to mock controls. Individual symbols represent independent biological replicates connected by dashed lines. N = 3 independent experiments.

**(B)** Quantification of cholesterol levels in GPMVs derived from APP-FL- and C99-expressing CHO cells under vehicle (circles) and +GSI pretreated conditions (squares) measured using the Amplex Red cholesterol assay. Values are expressed as fold change relative to mock controls. Individual symbols represent independent biological replicates connected by dashed lines. N = 3 independent experiments.

**(C)** Representative images of GPMVs from APP-FL and C99 variants under vehicle and +GSI conditions. GPMVs were labelled with a general lipid tracer (*blue*), the lipid-domain marker DiI (*magenta*), and APP/C99 species detected with 4G8 (*yellow*). N = 2 independent experiments. Scale bars, 5 µm.

**(D)** Representative images from an independent set of experiments highlighting C99 (4G8, *yellow*) localization relative to lipid domains (*magenta*) in GPMVs derived from $C99^{YENPTY}$-expressing cells under vehicle and GSI treatment (+GSI) conditions. N = 2 independent experiments. Scale bars, 5 µm. Increased clustering of $C99^{YENPTY}$ at discrete membrane regions is observed following γ-secretase inhibition.

Individual symbols represent independent biological replicates (N = 3). Statistical significance was determined using two-way ANOVA with post hoc multiple-comparison testing. * $p < 0.05$, ** $p < 0.01$, *** $p < 0.001$.

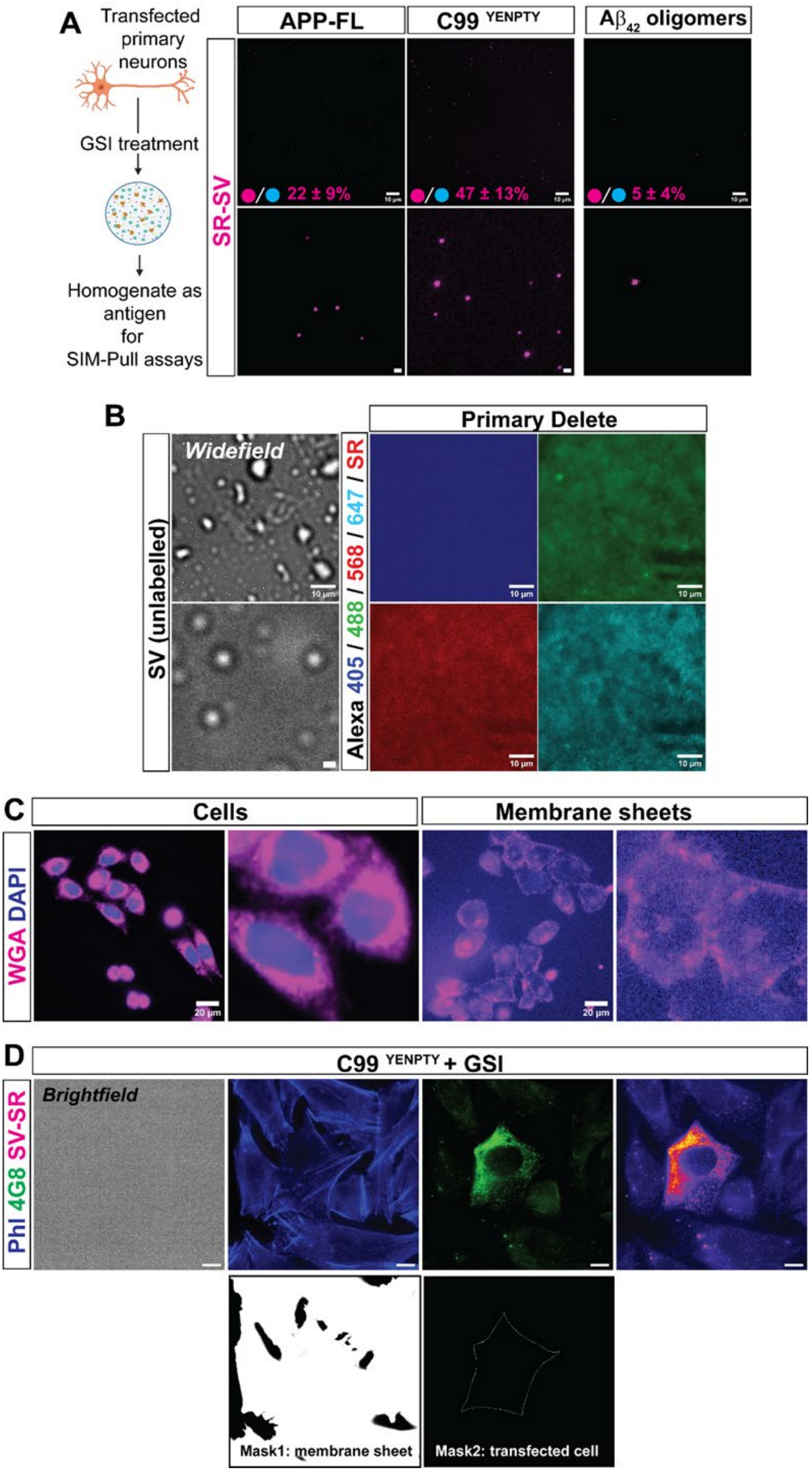


**Figure S6. Validation of synaptic vesicle tethering and membrane sheet analysis**

**(A)** Control SIMPull-based vesicle tethering assay using homogenates from primary neurons expressing APP-FL or C99[YENPTY] following GSI treatment. A$\beta_{42}$ oligomers were used as a negative control lacking the APP C-terminal domain. Values indicate the percentage of captured puncta associated with SynaptoRed-labelled synaptic vesicles, computed from N = 2 independent experiments. Scale bars, 10 µm; zoomed panels, 1 µm.

**(B)** Control images of unlabelled synaptic vesicle preparations imaged by widefield microscopy. Parallel fluorescence channels show minimal background signal under equimolar dye/secondary antibody conditions used for tethering assays. Scale bars, 10 µm; zoomed panels, 1 µm.

**(C)** Representative images of intact CHO cells and sonication-derived membrane sheets stained with WGA (magenta) and DAPI (blue). Intact cells show WGA-labelled plasma membranes surrounding DAPI-positive nuclei, whereas membrane sheets appear as flattened WGA-positive structures lacking nuclear signal, consistent with exposure of the cytosolic membrane face. Scale bars, 20 µm; zoomed panels, 5 µm.

**(D)** Representative membrane sheet–based vesicle tethering assay from GSI-treated $C99^{YENPTY}$-expressing CHO cells. Membrane sheets were labelled with phalloidin (*blue*), C99 species with 4G8 (*green*), and recruited SynaptoRed-labelled synaptic vesicles depicted in *magenta*. Lower panels show representative masks used for quantification: *mask 1*, total membrane sheet area generated from phalloidin; *mask 2*, APP/C99-positive transfected membrane area generated from 4G8 signals. Tethered SV-SR puncta were quantified in Fiji/ImageJ using particle analysis and expressed relative to membrane area. Scale bars, 5 µm.
